# Herbivore and carnivore microbial signatures indicate diet and health variation in human gut microbiota

**DOI:** 10.64898/2026.11.10.765789

**Authors:** Hiroto Miura, Tomohisa Takagi, Dai Ujihara, Akane Konno, Machi Oda, Shodai Ishikawa, Hiromi Ikeda, Kentaro Nagaoka, Hiroaki Kitae, Satoaki Matoba, Takamitsu Tsukahara, Yuji Naito, Ryo Inoue

## Abstract

Diet is a key factor shaping gut microbiota in mammals, with herbivores and carnivores harboring distinct microbial communities. However, despite humans being highly omnivorous, it remains unclear whether their gut microbiota more closely resembles that of herbivores or carnivores or how individual-level variations along this spectrum are associated with human health status. We compared gut microbiota profiles of 858 samples from 109 mammalian species and constructed a Herbivore-Carnivore (HC) index based on bacterial groups characteristic of each diet. The HC-index positioned humans between herbivores and carnivores and showed that traditional hunter-gatherers and nonhuman apes retained herbivore-leaning profiles, whereas urbanized humans exhibited a shift toward a carnivore-leaning state. By applying the HC-index to >72,000 participants across multiple cohorts worldwide, we found that elevated HC-index values were associated with plant-based and fiber-rich diets, greater microbial diversity, enhanced potential to degrade plant-derived carbohydrates, favorable clinical measures, and lower prevalence of self-reported disease. We demonstrated that the HC balance of gut microbiota captures a conserved diet-related gradient across mammals and provides an interpretable framework linking human diet, microbial function, and health.

## INTRODUCTION

The human gut microbiota plays a pivotal role in health/disease, and diet is one of the strongest factors shaping its composition and function^1–4^. The significance of diet is understood not only through the direct physiological effects of nutrients but also via indirect impacts mediated by the gut microbiota^3,4^. In this context, epidemiological and clinical evidence firmly establish the health benefits of dietary fiber; its higher intake has been associated with a reduced risk of diseases, including cardiovascular issues, type 2 diabetes, and colorectal cancer^3,5,6^. These benefits are largely attributed to the microbial fermentation of dietary fiber in the gut to produce bioactive metabolites, such as short-chain fatty acids (SCFAs)^7^. However, a fundamental question remains largely unexplained: despite humans being extremely omnivorous, why does dietary fiber exert such profound effects on our health, given that its benefits are largely microbiota-dependent and cannot be explained by host digestive capacity alone?

From an evolutionary perspective, this dependency may be rooted in our phylogenetic and dietary history. Most extant primates closely related to humans, such as chimpanzees and gorillas, are herbivores or frugivores^8^. In addition, paleoanthropological and archaeological evidence suggests that early hominins primarily consumed plant-based diets^9^. Although the evolution of the genus *Homo* involved progressive adaptations to meat consumption and developing sophisticated omnivory, plants remained the main dietary component^10,11^. Indeed, recent microbiological studies on paleofeces have revealed that the intestines of nonhuman primates and ancient humans harbor abundant cellulose-degrading bacteria^12^. Furthermore, the “Hadza,” who maintain a traditional hunter-gatherer lifestyle with plant-based diets, harbor gut microbiota with significantly enhanced fiber-degrading capacity compared to those of industrialized populations^13^. These observations suggest that the human gut microbiota may still retain “herbivorous traits,” which may explain why dietary fiber is crucial for maintaining health.

Across animals, it is well documented that gut microbial composition is driven more strongly by dietary habits^14,15^. The gut microbiota of herbivores is highly diverse and enriched in fiber-degrading bacteria, whereas that of carnivores exhibits lower diversity and a predominance of protein- and lipid-metabolizing bacteria^16,17^. Omnivores occupy an intermediate position, displaying compositional biases that reflect their primary diet^15,16^. Nevertheless, a pertinent question remains: does the gut microbiota of “highly omnivorous” modern humans more closely resemble that of herbivores or carnivores? Although studies have compared the human gut microbiota with those of several animal species, comprehensive research that positions the modern human gut microbiota along a continuous dietary spectrum from herbivory to carnivory is lacking. Moreover, none of these evaluated individual differences in the modern human gut microbiota along this herbivore-carnivore spectrum or investigated how these variations relate to host health status. These knowledge gaps could be addressed by a cross-sectional comparison of animal gut microbiomes, identifying bacterial taxa characteristic of carnivory or herbivory, and constructing a quantitative index based on their relative abundances.

The present study compares the gut microbiota of diverse animal species with distinct dietary patterns to construct a novel quantitative index that captures diet-driven differences in microbiota. Using this index, we position the gut microbiota of modern humans along the animal dietary spectrum. We then apply this metric to a large-scale human cohort to verify whether the index reflects variations in individual-level dietary patterns. Finally, we investigate the associations between this microbial index and host health outcomes, such as metabolic disease risk, thereby comprehensively elucidating the significance of “retained herbivorous traits” in maintaining human health.

## RESULTS

### Comparison of the human gut microbiota across diverse animal species

We analyzed 858 fecal microbiota samples from 109 mammalian species, comprising 35 carnivore species (*n* = 343 samples), 50 herbivores (*n* = 354 samples), and 24 omnivores (*n* = 161 samples) (Fig. 1a and Supplementary Table S1). Microbial alpha diversity differed significantly among dietary groups (*p* < 0.001); herbivores exhibited the highest diversity, followed by omnivores, with carnivores showing the lowest (Fig. 1b). Principal coordinate analysis based on beta diversity showed that microbial communities separated clearly along the PC1 axis (*R*^2^ = 0.24, *p* < 0.001, PERMANOVA) (Fig. 1c). Notably, omnivores were intermediate between herbivores and carnivores, reflecting a continuous dietary spectrum. To position humans within this range, we integrated fecal microbiota profiles from modern urbanized humans (Japanese) and traditional Hadza hunter-gatherers. The former clustered closer to carnivore-leaning omnivores, whereas the Hadza mapped within the herbivore cluster (Fig. 1d). While human populations with traditional dietary habits shared gut microbiota features with herbivores, the urbanized human microbiome shifted toward a carnivore-specific state.

**Fig. 1.**
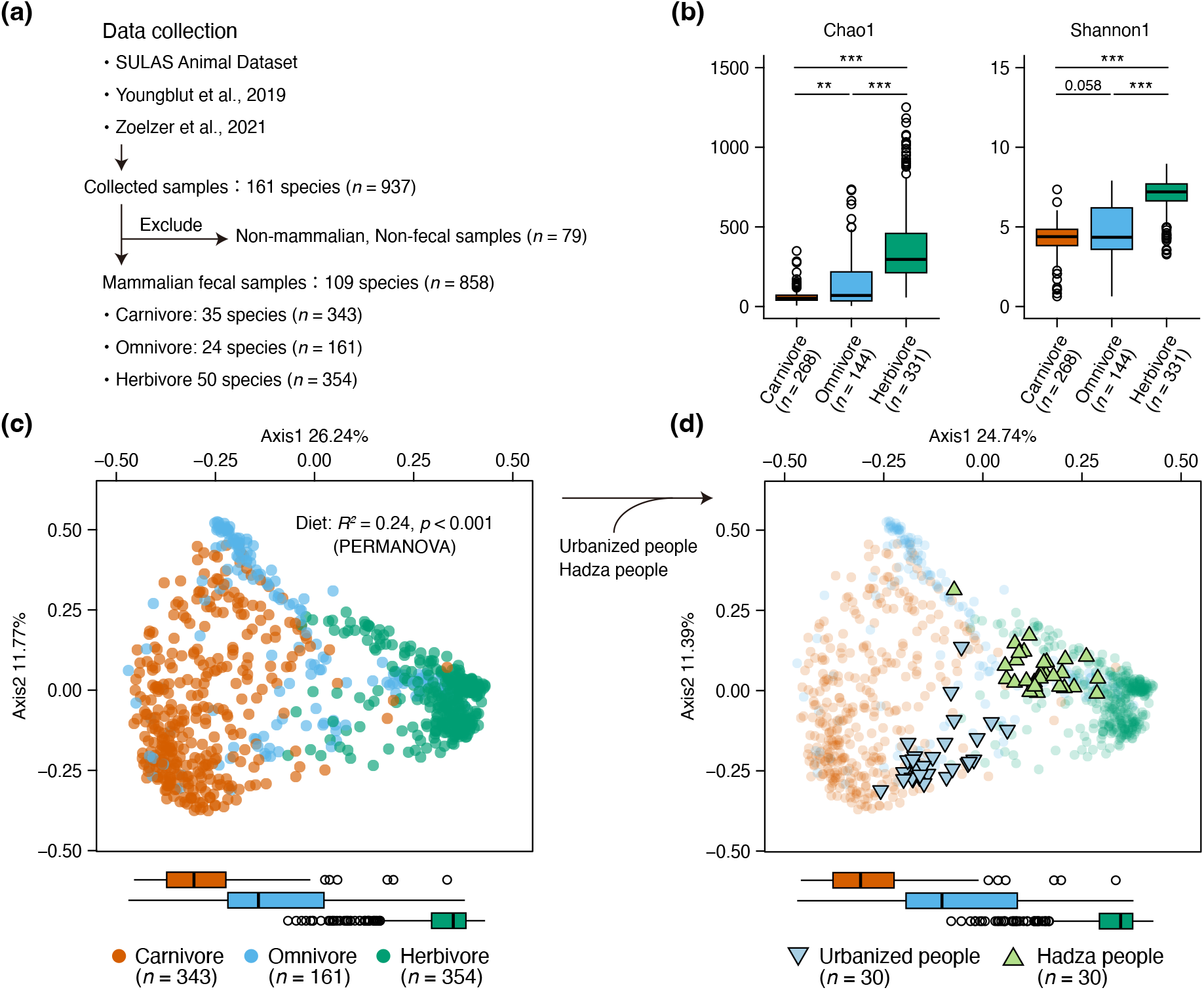
Comparison of gut microbiota profiles between humans and diverse animal species. (**a)** Schematic overview of the analytical workflow. Bioinformatic procedures are detailed in the Methods section. (**b**) Alpha diversity of gut microbiota across animal dietary groups. Alpha diversity indices were calculated after rarefying samples to 5,000 reads per sample; samples with < 5,000 reads were excluded. Statistical significance was assessed using a Kruskal–Wallis test followed by two-sided pairwise Wilcoxon rank-sum tests with Benjamini–Hochberg correction. Significance levels are indicated as **adjusted *p* < 0.01 and ***adjusted *p* < 0.001. (**c**) Principal coordinates analysis (PCoA) based on Bray-Curtis dissimilarities of gut microbial composition across diverse animal species. Box plots below the horizontal axis show the distribution of samples along PCoA axis 1. The proportion of variance explained by diet was assessed via a permutational multivariate analysis of variance (PERMANOVA) with 999 permutations. (**d**) PCoA based on Bray-Curtis dissimilarities of gut microbial composition comparing diverse animal species with the inclusion of urbanized people and Hadza people. Urbanized and Hadza samples were randomly subsampled from the original cohorts, with 30 samples included from each human population, as described in the Methods.

To quantitatively evaluate these diet-driven variations in gut microbiota, we developed an index based on the relative abundances of specific microbial taxa enriched in carnivores and herbivores (Fig. 2a). Among 963 microbial taxa detected in either of the two, we selected 61 candidates with a prevalence ≥ 50% in either group (Supplementary Table S2). Using indicator species analysis^18^, we then identified four carnivore- and 17 herbivore-associated taxa (*p* < 0.001 and indicator value > 0.85 for carnivore- or > 0.90 for herbivore-associated taxa; Supplementary Table S2). After excluding archaea and unclassified taxa at genus or genus-equivalent level, we finally identified four carnivore-associated taxa (C-taxa), including *Blautia* spp., *Escherichia–Shigella* spp., *Fusobacterium* spp., and *Peptoclostridium* spp., and 13 herbivore-associated taxa (H-taxa), comprising Bacteroidales p-251-o5, Christensenellaceae R-7 group, *Eubacterium coprostanoligenes* group, Oscillospiraceae NK4A214 group, Oscillospiraceae UCG-002, Oscillospiraceae UCG-005, Oscillospirales UCG-010, Prevotellaceae UCG-001, Prevotellaceae UCG-003, Prevotellaceae UCG-004, Rikenellaceae RC-9 group, *Ruminococcus* spp., and *Treponema* spp.

**Fig. 2.**
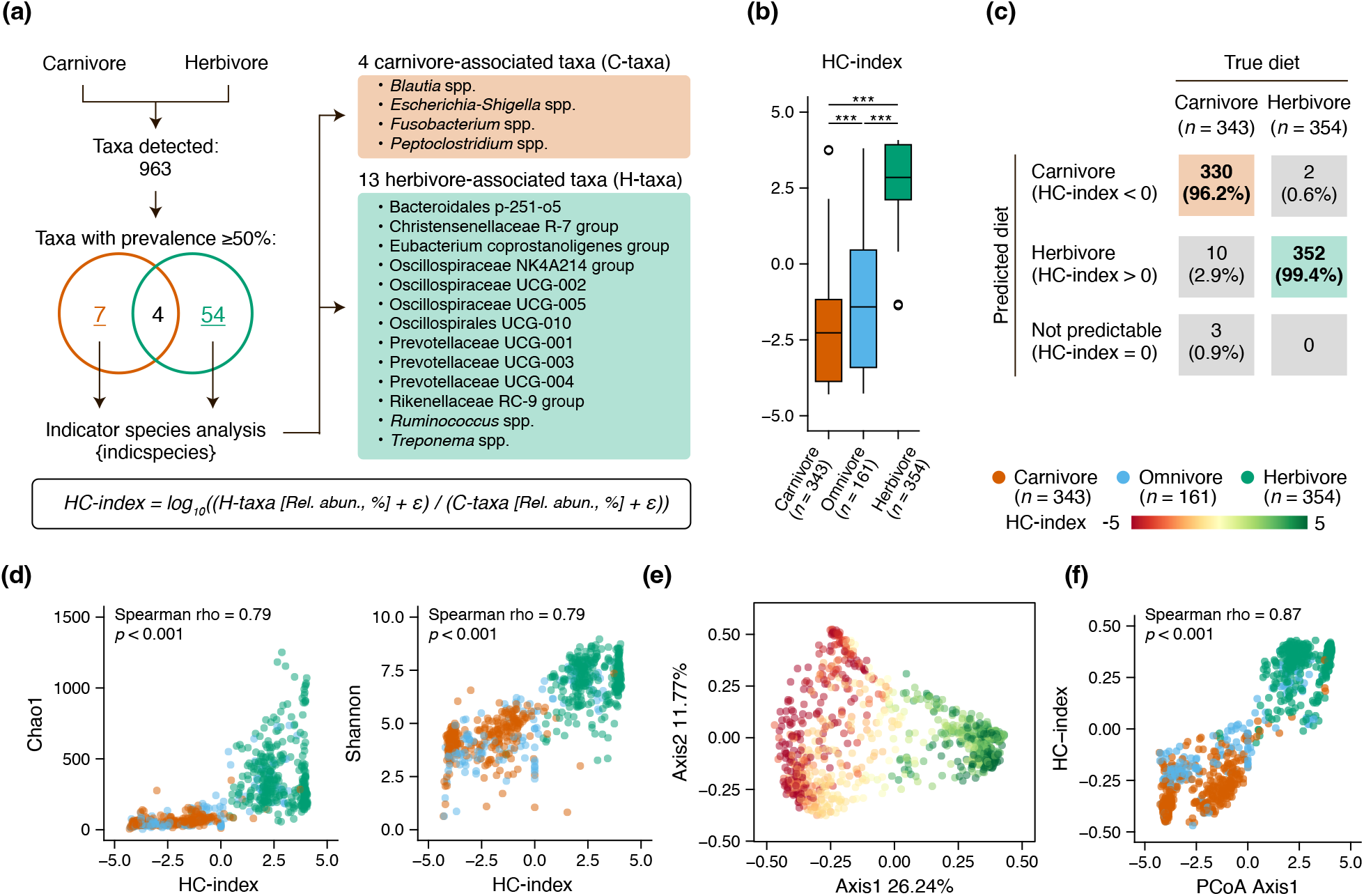
Development and validation of the herbivore-carnivore index. **(a**) Schematic overview of the analytical workflow used to construct the herbivore-carnivore (HC) index. Bioinformatic procedures are detailed in the Methods section. (**b**) Comparison of HC-index values across animal dietary groups. Statistical significance was assessed using a Kruskal–Wallis test followed by two-sided pairwise Wilcoxon rank-sum tests with Benjamini–Hochberg correction. Significance levels are indicated as ***adjusted *p* < 0.001. (**c**) Confusion matrix showing the performance of HC-index-based classification of carnivore and herbivore samples. Values indicate the number of samples, with percentages shown in parentheses. (**d**) Associations between the HC and alpha diversity indices, including Chao1 richness and Shannon diversity. Correlation coefficients and P values were calculated using two-sided Spearman’s rank correlation tests. (**e**) PCoA of Bray-Curtis dissimilarities across diverse animal species, with samples colored according to HC-index values. The ordination was identical to that shown in Fig. 1c. (**f**) Association between the HC-index and PCoA axis 1 coordinates presented in panel (e). Correlation coefficients and P values were calculated via a two-sided Spearman’s rank correlation test.

To express the compositional balance between H-taxa and C-taxa, we defined the Herbivore-Carnivore index (HC-index) as follows:

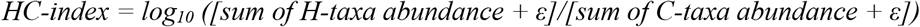

To account for variations in sequencing depth, the detection limit was set at 0.01% (assuming 1 read per 5,000–10,000 reads), and a pseudo-count (ε) of 0.005% (half of the detection limit) was added to handle zero abundances.

We calculated the HC-index of the animals used (Supplementary Table S3) and validated its applicability by comparing it across host diets and examining its correlation with overall microbial community structures. 1) The HC-index differed significantly (*p* < 0.001) among host dietary groups, with herbivores exhibiting the greatest values, followed by omnivores and carnivores (Fig. 2b). Furthermore, the index significantly correlated with the dietary proportion of plant-based foods reported in the EltonTraits database^19^ (Spearman rho = 0.76, *p* < 0.001; Supplementary Fig. S1); 2) Using an HC-index threshold of 0, we achieved high predictive accuracy, correctly classifying 96% and 99% of carnivore and herbivore samples, respectively (Fig. 2c); 3) The HC-index was strongly positively correlated with alpha diversity (Spearman rho = 0.79 for both Chao1 and Shannon, *p* < 0.001; Fig. 2d); and 4) beta diversity, specifically along the PC1 axis (Spearman rho = 0.87, *p* < 0.001; Fig. 2e and 2f). These correlations with alpha and beta diversity remained remarkable even when analyzed within individual dietary groups, indicating that the index captures the intra-diet gradient of microbial composition rather than merely reflecting group-level differences. These findings suggest that the HC-index is a useful tool for quantitatively evaluating diet-driven variations in gut microbial composition.

We calculated the HC-index for urbanized people, the Hadza people, and the ape groups most closely related to humans, using a larger sample size from an external dataset^20,21^. The Hadza and nonhuman apes maintained positive index values, whereas only urbanized humans showed negative values (Supplementary Fig. S2). Thus, while the human gut microbiota retains the potential to be within a herbivore-leaning state, urbanization has driven it toward a carnivore-leaning state.

### Association of the HC-index with dietary patterns in a human cohort

To validate the applicability of the HC-index in human gut microbiota, we calculated this metric based on the Human Microbiome Compendium dataset^22^, which aggregates gut microbiota profiles from multiple cohorts worldwide. After filtering samples based on metadata (see Methods), we analyzed a total of 38,832 adults across seven regions. While the distribution patterns of the HC-index values and the relative abundances of constituent taxa varied by region, inter-individual variations in the HC-index consistently spanned both positive and negative values across populations (Supplementary Figs. S3 and S4). Furthermore, the HC-index was markedly and positively correlated with microbial alpha diversity indices (Chao1 and Shannon) across all geographic regions (Supplementary Table S4). These results demonstrated that the HC-index serves as a robust metric across diverse human populations, confirming that while modern populations are carnivore-leaning on average, individual variations spanning the herbivore-carnivore spectrum exist globally.

Given these findings, we evaluated whether the HC-index reflects inter-individual differences in gut microbiota composition driven by individual dietary patterns. We utilized the Japanese Microbiome Integrated Cohort (J-MIC) cohort dataset, our largest original study group, comprising 26,783 adults who underwent standardized microbial analysis and provided comprehensive questionnaire-based background data. The J-MIC is a large-scale, real-world gut microbiome cohort established based on data accumulated through Flora Scan®, a commercially available gut microbiome testing service available in Japan. In this cohort, the HC-index was widely distributed, spanning positive and negative values (*n* = 4,416 for HC-index ≥ 0, *n* = 22,367 for HC-index < 0; IQR = −2.42 to −0.24; Fig. 3a), similar to that observed in the global dataset for Southeast Asia. Age- and sex-adjusted multinomial logistic regression analyses revealed remarkable associations between the HC-index and specific dietary habits. Individuals with a higher HC-index showed a prominent preference for high-fiber diets, characterized by the frequent consumption of green and yellow vegetables and fermented foods, whereas the index was negatively associated with the regularity of pork and beef consumption (Fig. 3b). Individuals with an enhanced HC-index exhibited not only a higher prevalence of dietary fiber intake but also greater awareness of dietary fiber, including a stronger intention to consume fiber-rich foods (Fig. 3c). This relationship between HC-index and dietary habits was validated using the American Gut Project dataset^23^, as an independent cohort from another country. Similar to our J-MIC cohort, higher HC-index scores were associated with more frequent consumption of vegetables and fruits, but not with meat intake (Supplementary Fig. S5). Finally, we quantitatively validated these relations employing another independent group, the Kyotango cohort^24^, which comprised 770 adults with detailed dietary intake records derived from a brief diet history questionnaire (BDHQ)^25^ and passed through the same analytical pipeline as the J-MIC cohort. Analyses of age- and sex-adjusted correlations confirmed that the HC-index associated positively with absolute intakes of nutrients abundant in vegetables and fruits, including dietary fiber, carotenoids, and vitamins (Fig. 4 and Supplementary Table S5). Energy-adjusted intakes of water-soluble, insoluble, and total dietary fiber differed significantly across HC-index quartiles, with higher HC-index quartiles showing greater fiber consumption (Supplementary Fig. S6). In contrast, statistically significant associations were not observed between the HC-index and meat consumption. The findings across multiple independent cohorts robustly confirm that the HC-index closely reflects individual variations in gut microbiota composition driven by dietary patterns, particularly plant-based foods, thereby capturing herbivore- and carnivore-leaning states.

**Fig. 3.**
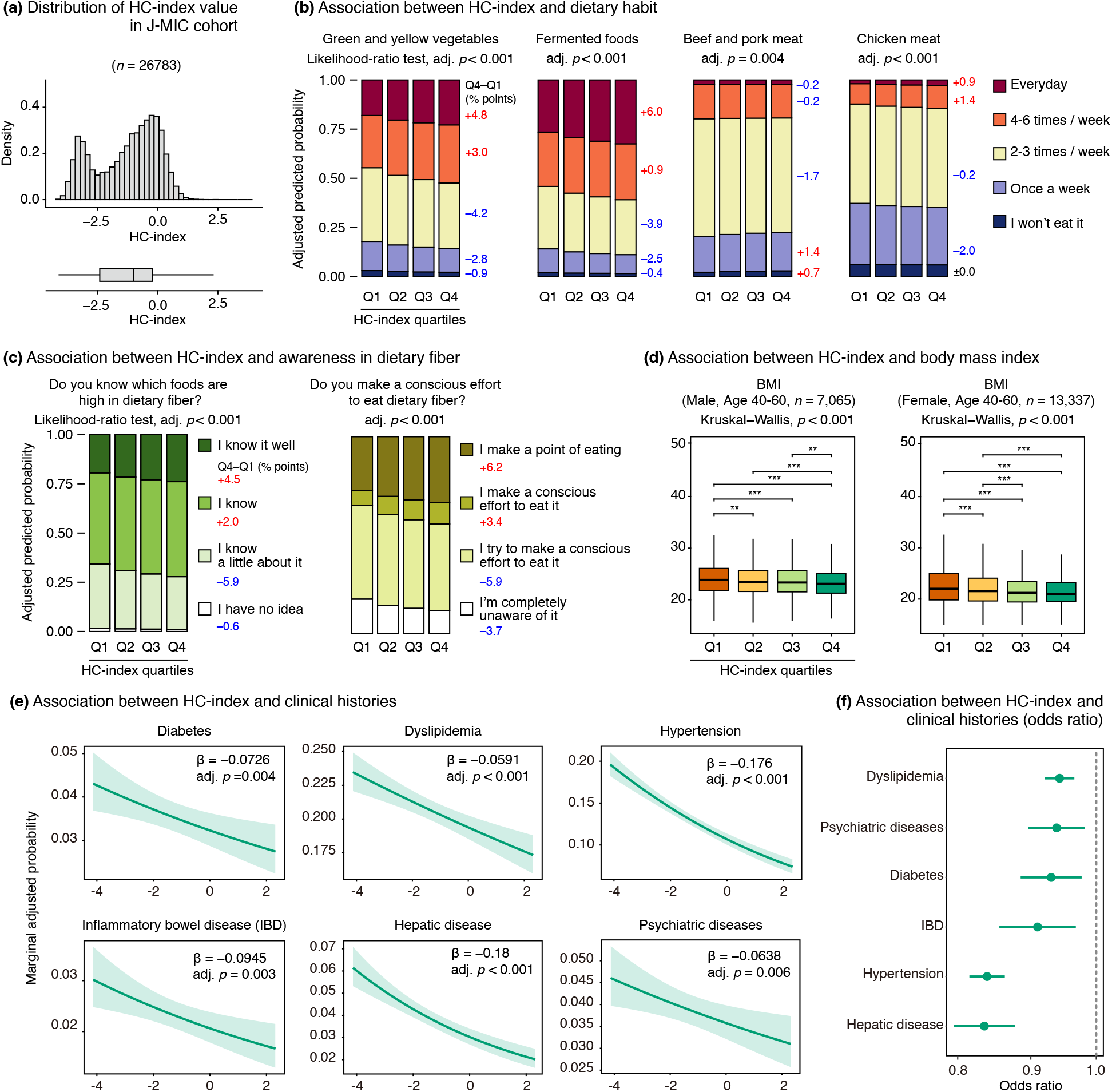
Association between the HC-index and dietary habits, body mass index, or clinical histories in the J-MIC cohort. (**a**) Distribution of HC-index values in the J-MIC cohort (*n* = 26,783). (**b**) Associations between HC-index and the questionnaire-derived frequency of intake for selected food items. Age- and sex-adjusted multinomial logistic regression models were used to predict marginal probabilities for each response category, shown as stacked bars across HC-index quartiles. Statistical significance of the HC-index was evaluated using a likelihood-ratio test comparing a full model including the HC-index, age, and sex with a reduced model comprising only age and sex. P values were adjusted for multiple testing by applying the Benjamini–Hochberg procedure. For interpretation, Q4−Q1 differences in adjusted predicted probabilities are shown next to the stacked bars in percentage points; positive values indicate a higher predicted probability in Q4 (higher HC-index), whereas negative values indicate a higher predicted probability in Q1 (lower HC-index). The corresponding model-based curves are presented in Supplementary Fig. S7. (**c**) Association between HC-index and awareness of dietary fiber intake. Analyses and data presentation were performed as described in panel (b). **(d)** Comparison of body mass index (BMI) across HC-index quartiles. Data were stratified by sex and restricted to participants aged 40–60 years. Statistical significance was assessed utilizing a Kruskal–Wallis test followed by two-sided pairwise Wilcoxon rank-sum tests with Benjamini–Hochberg correction. Significance levels are indicated as *adjusted *p* < 0.05, **adjusted *p* < 0.01 and ***adjusted *p* < 0.001. (**e**) Age- and sex-adjusted associations between HC-index and clinical histories were estimated via binary logistic regression models. The green line and shaded area indicate the fitted regression line and its 95% confidence interval, respectively. Adjusted *β* denotes the unstandardized regression coefficient post-adjustment for age and sex. Statistical significance was assessed employing likelihood-ratio tests with Benjamini– Hochberg correction. (**f)** Age- and sex-adjusted odds ratios for associations between HC-index and clinical histories. Odds ratios are shown per one-unit increase in HC-index. Points indicate odds ratios, and whiskers represent 95% confidence intervals.

**Fig. 4.**
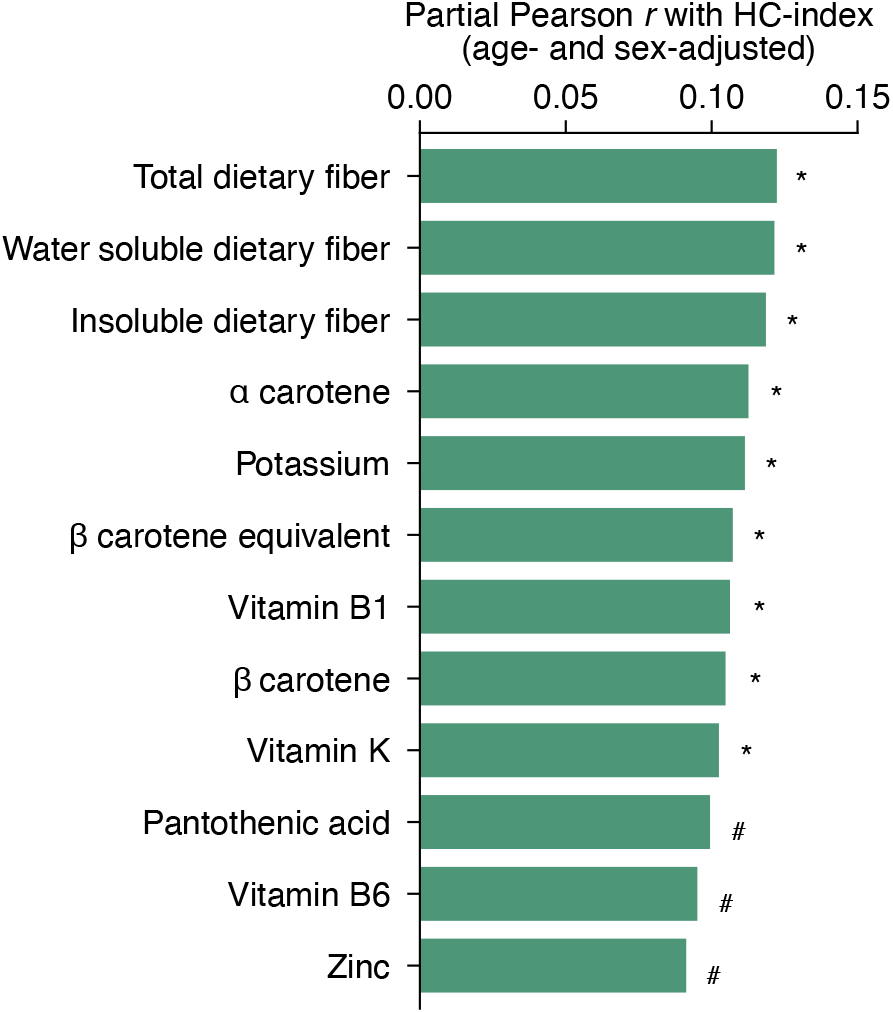
Correlation between the HC-index and nutrient intake in the Kyotango cohort. Age- and sex-adjusted associations between HC-index and absolute nutrient intake were assessed in the Kyotango cohort (*n* = 770). Nutrient intake was estimated as absolute daily intake using the Brief-type self-administered Diet History Questionnaire. Partial Pearson’s correlation coefficients and corresponding P values were calculated post-adjustment for age and sex. P values were corrected for multiple testing using the Benjamini–Hochberg procedure. Only nutrients with adjusted *p* < 0.10 are shown. Significance levels are indicated as *adjusted *p* < 0.05, ^#^adjusted *p* < 0.10. Complete results are provided in Supplementary Table S5.

### Association of the HC-index with human physiological and clinical status

Having established the HC-index as an indicator of diet-driven gut microbial patterns, we characterized its association with human physiological and clinical status. Using the J-MIC cohort, an age- and sex-adjusted analysis revealed that the HC-index was negatively correlated with body mass index (Fig. 3d), associating a higher HC-index profile with a leaner phenotype. Furthermore, a higher HC-index was associated with a decreased prevalence of several clinical histories, including hepatic disease, hypertension, inflammatory bowel disease, diabetes, psychiatric disease, and dyslipidemia (Figs. 3e and 3f). Thus, the HC-index reflects not only diet-driven microbial variations but also likely the associated physiological and clinical states.

The clinical relevance of the HC-index was evaluated employing the KPUM dataset^26^, which comprised 1,803 adults with comprehensive clinical blood test records and underwent the same analytical pipeline as the J-MIC cohort. We observed marked correlations between the HC-index and biochemical parameters reflecting liver and kidney function and lipid metabolism (Fig. 5 and Supplementary Table S6). Enhanced HC-index values correlated negatively with serum levels of aspartate aminotransferase (AST), alanine aminotransferase (ALT), and triglycerides (TGs), and positively with urine pH and blood urea nitrogen. These clinical data demonstrate that an herbivore-leaning gut microbiota profile is closely related to improved liver and kidney function and systemic lipid metabolism.

**Fig. 5.**
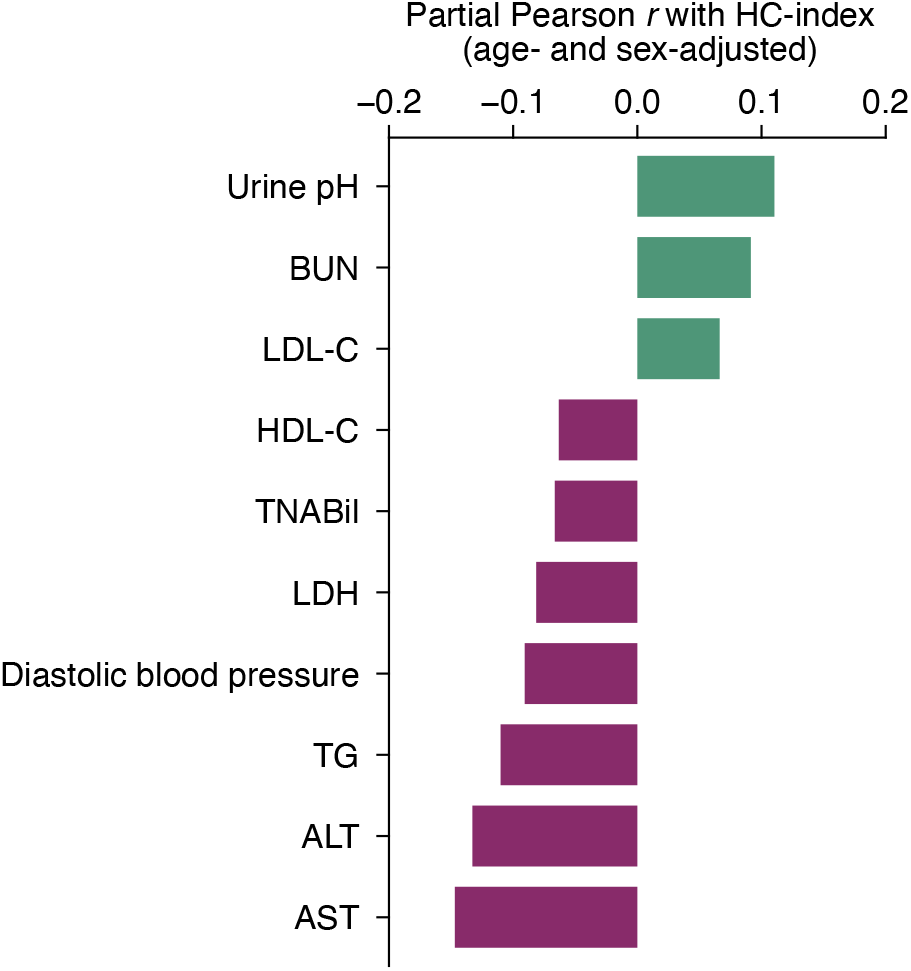
Correlation between HC-index and blood biochemical parameters in the KPUM dataset. Age- and sex-adjusted associations between HC-index and blood biochemical parameters were assessed in the KPUM dataset (*n* = 1,803). Partial Pearson’s correlation coefficients and corresponding P values were calculated post-adjustment for age and sex. P values were corrected for multiple testing using the Benjamini–Hochberg procedure. Only parameters with adjusted *p* < 0.05 are shown. Complete results are provided in Supplementary Table S6.

### Association of the HC-index with the functional potential of gut microbiota

We investigated the link between HC-index and the functional potential of gut microbiota using shotgun metagenomic data derived from 156 adults belonging to the J-MIC cohort. The overall profile of functional genes identified based on KEGG annotation exhibited a gradual shift corresponding to the HC-index (Fig. 6a), and genes recognized via CAZyme annotation showed a clearer separation of individuals (Fig. 6b). To focus on carbohydrate metabolism potential, we then compared the abundance of CAZyme genes related to carbohydrate degradation between individuals in the lowest and highest HC-index quartiles, Q1 and Q4, respectively. Linear discriminant analysis with Wilcoxon rank-sum tests revealed that multiple CAZymes involved in plant-derived carbohydrate degradation were markedly more abundant in Q4, whereas those associated with animal-derived carbohydrate and host glycan breakdown were remarkably predominant in Q1 (Fig. 6c).

**Fig. 6.**
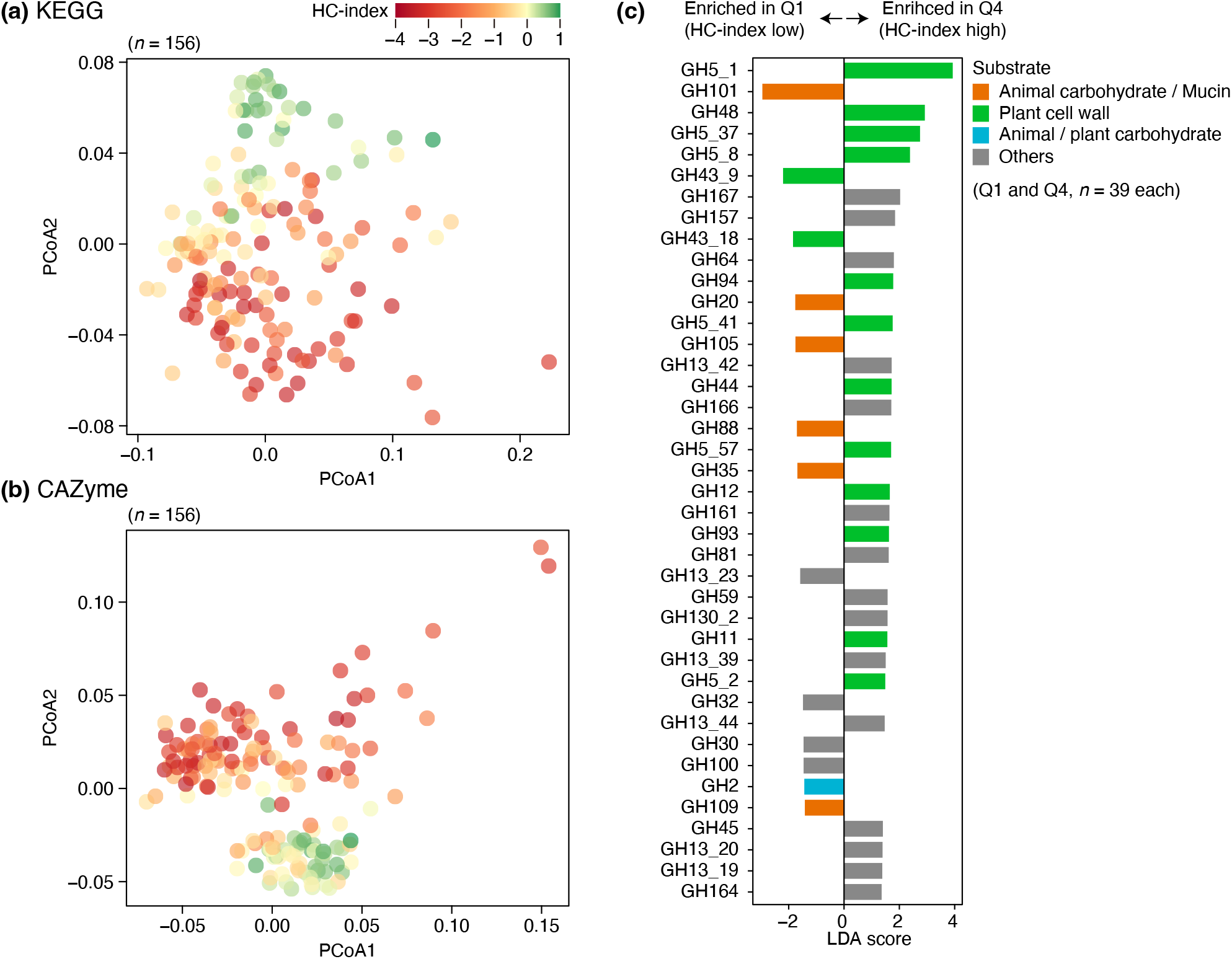
Associations between HC-index and metagenomic functional profiles in the J-MIC cohort. Principal coordinates analysis (PCoA) of Bray-Curtis dissimilarities via **(a)** KEGG-based functional gene profiles and **(b)** CAZyme gene profiles. Bray-Curtis dissimilarities across samples were calculated based on TPM-normalized gene abundances. Individual plots are colored according to HC-index values. **(c)** Family-based glycoside hydrolase (GH) genes enriched in subjects with lower and higher HC-index values. TPM-based abundance of GH genes was compared between the lowest HC and highest HC-index quartiles (Q1 and Q4, respectively) using two-sided Wilcoxon rank-sum tests with Benjamini– Hochberg correction. Only GH genes with adjusted *p* < 0.05 are presented. Higher and lower LDA scores indicate genes enriched in subjects with higher and lower HC indices, respectively. GH families were categorized by predicted substrate class—targeting plant cell wall carbohydrates, animal-derived carbohydrates, mucin-related glycans, or other substrates— based on a previous study (Smits et al., 2017).

### Changes in HC-index with diet intervention relate to microbial diversity and fermentation

To examine whether the HC-index can be altered by dietary intervention, we analyzed data from our previous study involving randomized controlled dietary intervention^27^. There, healthy adults received dietary fiber supplementation for 4 weeks at either a low (2.2 g/day; *n* = 55) or a high (8.2 g/day; *n* = 55) dose. After calculating the HC-index, we evaluated baseline-adjusted changes in the index during the intervention period (Fig. 7a). In the high-fiber group, this change was significantly positive from weeks 0 to 2 (*p* = 0.035), but was insignificant in the low-fiber group. The mean change in HC-index from weeks 0 to 4 was also positive in the high-fiber group, although it did not reach statistical significance (*p* = 0.287). These findings suggest that the HC-index may be responsive to dietary intervention and that dietary fiber supplementation may increase the HC-index, although further validation is required.

**Fig. 7.**
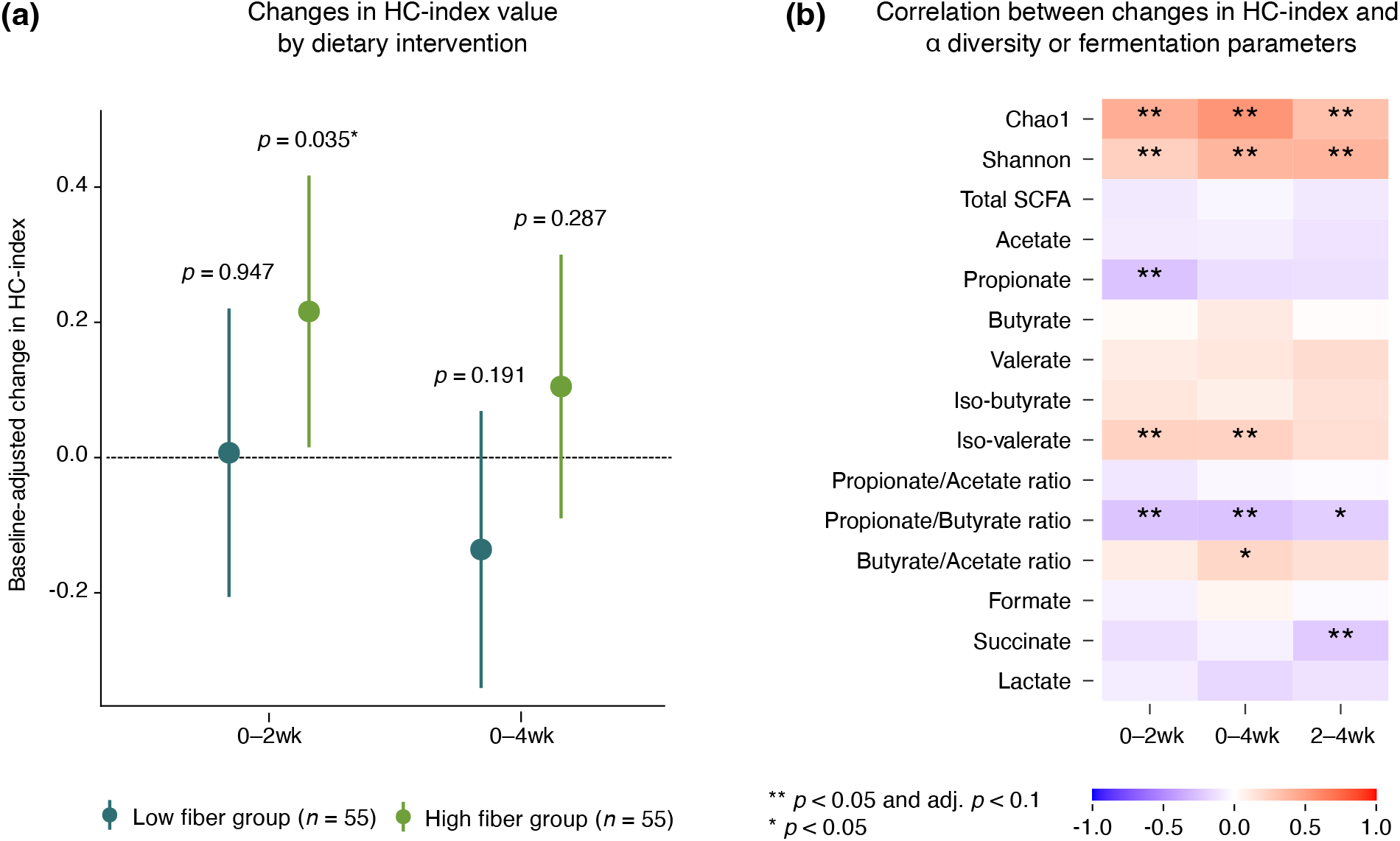
Changes in HC-index by dietary intervention and their relation to microbial alpha diversity and fermentation parameters. Healthy adults received dietary fiber supplementation for 4 weeks at either a low dose (2.2 g/day; *n* = 55) or a high-dose (8.2 g/day; *n* = 55), and fecal microbiota and fermentation parameters were analyzed at 0, 2, and 4 weeks of the trial**. (a)** Baseline-adjusted changes in the HC-index during the intervention period. Points indicate estimated baseline-adjusted changes in the HC-index, and whiskers indicate 95% confidence intervals. **(b)** Correlations between changes in HC-index, microbial alpha diversity, and fermentation parameters with dietary fiber supplementation. The heatmap shows Pearson’s correlation coefficient values (red, positive; blue, negative). P values were calculated and adjusted using the Benjamini–Hochberg procedure. Significance levels are indicated as * nominal *p* < 0.05, ** nominal *p* < 0.05 with adjusted *p* < 0.10.

We next examined whether intervention-induced changes in HC-index were associated with changes in microbial alpha diversity and fecal fermentation parameters (Fig. 7b). HC-index alterations were markedly and directly correlated with alpha diversity changes. We also observed significant associations between HC-index changes and fecal SCFA profiles. Across all intervention intervals examined—weeks 0–2, 0–4, and 2– 4—HC-index alterations showed consistent, significant inverse correlations with changes in the propionate– butyrate ratio. As fecal SCFA concentrations reflect the balance between microbial production, host absorption, and fecal excretion, they cannot be directly interpreted as absolute SCFA production. Nevertheless, these results suggest that HC-index changes may be associated with shifts in microbial fermentation balance during dietary intervention.

### Comparison of explanatory performance between HC and alpha diversity indices

Across all cohorts, HC-index was strongly and positively correlated with alpha diversity indices (Supplementary Table S4). To investigate whether the HC-index captured varied information from alpha diversity, explanatory performance was compared between the HC and alpha diversity indices utilizing the J-MIC cohort dataset (Fig. 8). For categorical variables, in addition to age and sex, likelihood-ratio statistics from models including the HC or alpha diversity indices were compared. For numerical variables, the strength of association was compared using Spearman’s rho. HC-index showed larger likelihood-ratio statistics or more robust correlations than alpha diversity indices for the above-mentioned lifestyle- and disease-related variables, suggesting that the HC-index may capture relevant biological information not fully reflected by conventional alpha diversity metrics such as richness or evenness.

**Fig. 8.**
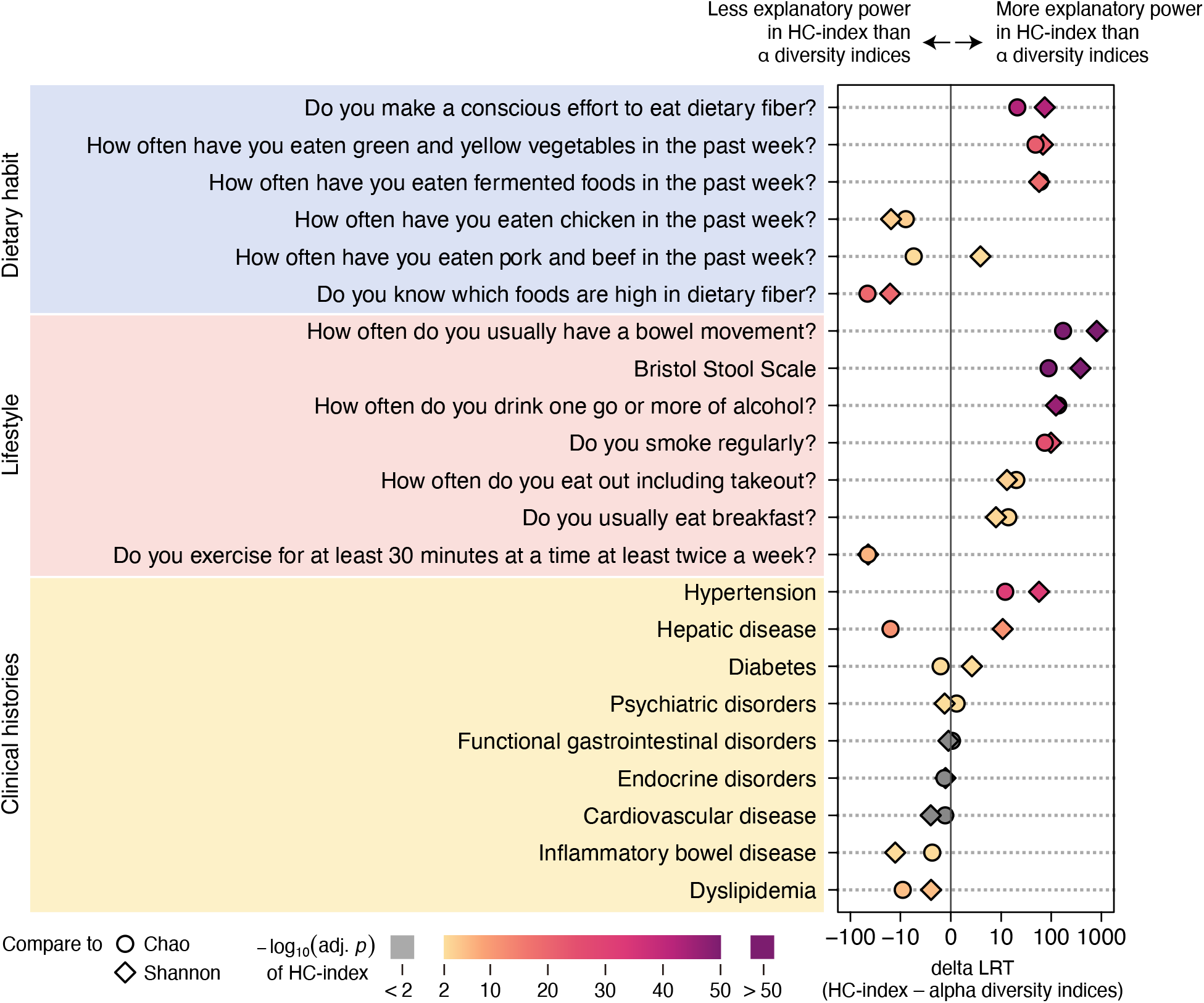
Comparison of the explanatory power of the HC and alpha diversity indices in the J-MIC cohort. Age- and sex-adjusted regression models were used to compare the explanatory power of the HC-index with that of alpha diversity indices (Chao1 and Shannon) for questionnaire-based variables related to dietary habits, lifestyle behaviors, and clinical histories in the J-MIC cohort. Multinomial logistic regression models were employed for categorical questionnaire variables, and binary logistic regression models were utilized for clinical histories. Model fit was evaluated via likelihood-ratio test statistics. Differences in likelihood-ratio test statistics were calculated as ΔLRT = LRT(HC-index) − LRT(alpha diversity index), where alpha diversity was represented by either Chao1 richness or Shannon diversity. Positive ΔLRT values indicate greater explanatory power of the HC-index than the corresponding alpha diversity index, while negative values suggest the contrary. Statistical significance was assessed employing likelihood-ratio tests, and P values were adjusted for multiple testing using the Benjamini–Hochberg procedure.

## DISCUSSION

In this study, we developed the HC-index to quantitatively evaluate the herbivore- and carnivore-leaning states of the animal gut microbiota. Although several studies have compared the gut microbiota of carnivorous and herbivorous animals^28^, this is the first report to translate these differences into a comprehensive and continuous index; we used this index to determine the position of modern human gut microbiota within a continuous mammalian dietary spectrum. While urbanized humans were mapped as carnivore-leaning omnivores, nonhuman apes and traditional Hadza hunter-gatherers distinctly clustered within the herbivore spectrum. These results demonstrate that although the human gut microbiota inherently retains a herbivore-typical potential, modern urbanized/industrialized lifestyles, including dietary shifts, have driven it toward a carnivore-type state, as noted by Makki et al.^5^ and Moraïs et al.^12^ As an example similar to the Hadza population, children from rural villages in Burkina Faso who primarily consume a plant-based diet harbored a distinct gut microbiota enriched in dietary fiber-degrading bacteria compared with urban individuals^29^; additionally, when the rural individuals migrate to urban environments, their gut microbiota shifted toward a more urbanized microbial composition^30–32^. These observations underscore the substantial impact of habitual diet, independent of host genetics, suggesting that an herbivore- or carnivore-characteristic gut microbiota is primarily shaped by environmental factors, including diet, rather than host genetic backgrounds^33^. These insights suggest that modern people are losing the herbivorous trait of gut microbiota due to urbanization/industrialization, including chronic dietary fiber deficiency^34,35^. Researchers estimate that our ancestors consumed more than 100 g of dietary fiber per day^36^, and the daily fiber intake of the Hadza ranges from 80 to 150 g^37^. Therefore, for modern urbanized humans, even fiber intakes that meet current dietary recommendations (e.g., approximately 20 g/day in Japan, 30 g/day in the US, and 25 g/day in Europe) may still be far below the amounts required to maintain a herbivore-leaning state by the gut microbiota. That is, under modern urbanized/industrialized lifestyles, the gut microbiota exists in a state of constant dietary fiber deficiency, which offers a mechanistic explanation for why fiber intake is so fundamentally vital for human health.

By applying the HC-index to modern human populations globally, we validated that the gut microbiota of modern humans has, on average, indeed shifted toward a carnivore-leaning state, irrespective of region. However, individual variations in the HC-index values spanned from negative to positive across all global populations examined, suggesting the presence of a continuous herbivore-carnivore spectrum even within modern humans. We also observed variations in the distribution patterns of the HC-index across regions and speculate that they are attributable to localized environmental factors, particularly regional lifestyles and cultural dietary habits^32,38^.

To elucidate the specific factors explaining such individual variation in HC-index, we analyzed data from multiple human cohort studies. Across these independent cohorts—the J-MIC, American Gut Project, and Kyotango datasets—the association between HC-index and habitual dietary patterns was robustly confirmed. Notably, enhanced HC-index values were consistently associated with a preference for plant-based foods, such as green and yellow vegetables and fruits, whereas lower scores were associated with higher meat consumption. This pattern demonstrates that the HC-index serves as an effective metric even within human populations, successfully capturing individual variations in gut microbial states driven largely by the intake of plant- and meat-based diets, aligning with the concept of the herbivore-carnivore spectrum. In addition, quantitative analysis of the Kyotango cohort demonstrated that the HC-index more prominently and continuously correlated with dietary fiber than with meat consumption. The greater impact of dietary fiber than meat intake can be explained by our hypothesis that the modern gut microbiota exists in a constant state of dietary fiber deficiency.

Beyond direct dietary choices, we observed that the HC-index also captured broader lifestyle behaviors. Specifically, elevated HC-index values were significantly associated with healthy lifestyle patterns, including regular exercise and habitual breakfast consumption, as well as lower frequencies of dining out, alcohol consumption, and smoking (Supplementary Fig. S7). These behaviors contrast with the typical features of urbanized/industrialized lifestyles, indicating that individuals with an herbivore-leaning gut microbiota tend to maintain a relatively traditional lifestyle. This finding is consistent with a recent large-scale digital cohort study reported by Singh et al.^39^, which demonstrated that unhealthy habits, particularly smoking and frequent alcohol intake, negatively correlate with both diet quality (e.g., the Healthy Eating Index) and gut microbial alpha diversity, the enhanced levels of which are well-known features of herbivorous microbiomes^3^. Therefore, the herbivore-leaning gut microbiota in modern humans is shaped by a traditional-leaning lifestyle centered on a plant-based diet.

In this study, we also examined the associations between HC-index and bowel movement frequency or stool consistency. Individuals with a higher HC-index had less frequent bowel movements and harder stool consistency (Supplementary Fig. S8). These findings may partly reflect the influence of a vegetable-rich diet, which could improve stool firmness and prolong intestinal transit time^39,40^. Although these results might suggest a constipation-like phenotype among individuals with a greater HC-index, such an interpretation should be cautious and requires further investigation^41–43^. Notably, Singh et al.^39^ also reported a negative association between alpha diversity and bowel movement frequency, while alpha diversity was positively associated with self-reported improved stool quality in this study. Future studies must further clarify these relationships by directly measuring fecal water content, together with other dietary and lifestyle factors that may influence bowel habits.

Importantly, diet- and lifestyle-driven variations in HC-index were associated not only with taxonomic shifts but also with functional changes in the gut microbiota, which may have implications for host health. Our metagenomic analyses revealed an enrichment of genes involved in the degradation of plant-derived carbohydrates in the higher HC-index group, suggesting microbial adaptation to fiber-rich diets; these diets are well established to enhance microbial SCFA production, particularly butyrate^5,44,45^. Individuals with a greater HC-index may harbor a gut microbial ecosystem characterized by a more butyrate-oriented fermentation profile.

This interpretation is further supported by our re-analysis of an earlier dietary intervention study^27^, in which high-dose dietary fiber supplementation increased the HC-index during the early phase of the intervention. Intervention-induced changes in the HC-index were consistently associated with changes in fecal SCFA profiles, particularly a decline in the propionate–butyrate ratio. As fecal SCFA concentrations reflect the net balance among microbial production, host absorption, and fecal excretion^46^, these results cannot be directly interpreted as alterations in absolute SCFA production. Although validation is required, they suggest that a higher HC-index is associated with a more butyrate-enriched fermentation pattern. Thus, the HC-index is not merely a static taxonomic marker but a diet-responsive microbiota index associated with shifts in microbial fermentation balance.

Butyrate plays a crucial role in maintaining the intestinal barrier, immune homeostasis, and metabolic regulation^3,5^. A greater HC-index may reflect a gut microbial environment that is more favorable for supporting host health. In contrast, the enrichment of genes for host glycan and animal-derived carbohydrate degradation in the lower HC-index group indicates a metabolic shift caused by a lack of dietary fiber. Under fiber-deficient conditions, the microbiota is forced to use alternative energy sources, including proteins and the host mucin layer^47–50^. This observation is supported by human dietary studies showing that low-carbohydrate, high-protein diets significantly suppress SCFA production and increase harmful products from protein fermentation, such as ammonia, phenolics, and hydrogen sulfide^51,52^. Therefore, an herbivore-leaning gut microbiota, represented by a higher HC-index, likely contributes to host health by maintaining SCFA production and suppressing potentially toxic protein fermentation.

These functional and fermentation shifts can explain the association between HC-index and host clinical phenotypes observed in this study. Individuals with a higher HC-index had fewer medical histories and an overall better health status than those with a lower index. Previous microbiome-derived indices, such as the Gut Microbiome–Health Index^53^ and Gut Microbiome Wellness Index 2^54^, suggest that composite microbial signatures can discriminate health status among humans. However, these indices were explicitly constructed employing human cohorts labeled according to health or disease status, making health status discrimination an intended design property. In contrast, the HC-index was derived independently of human health information, based solely on the relationship between host dietary ecology and gut microbial composition across nonhuman mammals, and was not optimized for any human dietary or clinical outcome. Notably, despite this phenomenon, the HC-index captured variation not only in human dietary patterns but also in lifestyle, health status, and disease-related phenotypes. This outcome represents a key conceptual novelty of the HC-index: rather than being designed as a classifier of health and disease status, it should be developed as a diet-associated ecological axis of the gut microbiome that can independently reveal health-relevant variations in humans.

Consistent with these broader associations with health, a higher HC-index was quantitatively associated with favorable clinical blood parameters related to liver function, kidney function, and lipid metabolism, which could also be explained from an evolutionary perspective. Selective constraints on metabolism-related genes expressed in the liver and kidneys, which play a central role in the metabolism of animal-derived nutrients such as proteins and lipids, were eased in correlation with herbivory, reducing metabolic capacity^55^. Modern humans who retain the traits of a herbivore-type metabolism may experience excessive metabolic stress when consuming habitually elevated amounts of animal-derived proteins and lipids. In such conditions, ammonia and urea detoxification may become overloaded, resulting in chronic stress on liver and kidney function^56^. Although causal relationships cannot be definitively proven within the current scope, our findings suggest that individuals with a more herbivore-leaning gut microbiota and who consume a diet rich in plant-based foods, as indicated by a high HC-index, maintain better overall physiological health.

In terms of the relationship between dietary habits and gut microbiota along the herbivore-carnivore axis, changes in alpha diversity are widely recognized^17,28^. In fact, the HC-index was significantly positively correlated with alpha diversity across all cohorts in this study, suggesting that it shares features with alpha diversity. However, when focusing on the fundamental causal relationship among diet, alpha diversity, and HC-index, the HC-index captured more information than alpha diversity, rather than merely reflecting it. That is, instead of a high-diversity state enhancing HC-index, a greater HC-index—which means a herbivore-leaning gut microbial environment—likely creates the ecological niches necessary to support and sustain an extremely diverse microbial community. The HC-index is more closely and fundamentally tied to dietary patterns than conventional alpha diversity indices. This assumption is strongly supported by our statistical validations, in which the HC-index consistently yielded markedly higher explanatory power (ΔR2 and ΔLRT) for dietary patterns and clinical disease associations than alpha diversity indices. These data indicate that to understand and predict host phenotypes and metabolic conditions, the structural balance along the herbivore-carnivore axis (the quality of composition) is more informative and practical than simply measuring the overall richness and evenness of the gut microbiota.

The HC-index provides a significant advantage in analytical robustness over alpha diversity and the relative abundance of specific taxa. The relative abundance of individual bacterial taxa fluctuates readily in response to biological and experimental noise, minor environmental changes, and personal factors. Given that the taxa constituting the HC-index vary across regions (Supplementary Fig. S4), a multi-taxon approach is essential to maintain global applicability. Consistent with this ecological stability, the HC-index correlated more remarkably with microbial alpha diversity than with any individual constituent taxon (Supplementary Table S7).

Furthermore, the HC-index is broadly applicable and practically accessible. Unlike proposed gut microbiome–health indices, which rely on shotgun metagenomics and relatively large sets of microbial features—50 species and 95 taxa for the Gut Microbiome–Health Index^53^ and the Gut Microbiome Wellness Index 2^54^, respectively—the HC-index was calculated based only on 17 taxa and can be derived from 16S rRNA gene sequencing data. In the case of a Japanese cohort, the descriptive and predictive accuracy of the HC-index was largely maintained even when the number of H- and C-taxa was narrowed down to a maximum of seven taxa (see Methods; Supplementary Fig. S9). Thus, compatibility with 16S rRNA gene sequencing may lower the technical and analytical barriers to implementation, while the small number of required taxa and the possibility of further panel reduction may enable rapid and simple targeted quantification approaches, such as qPCR, thereby facilitating broader application of the HC-index in the future.

Several limitations should be considered. First, our analyses relied on cross-sectional cohorts; thus, we cannot definitively establish a direct, temporal causal relationship among changes in HC-index, dietary patterns, and health status. Second, despite adjustment for age and sex, clinical history models may remain confounded by medication use, comorbidities, and lifestyle factors. Third, genus-level 16S rRNA gene profiling cannot resolve species- or strain-specific variations; therefore, HC-index taxa should be interpreted as ecological markers rather than functionally homogeneous organisms. Further longitudinal and interventional studies are warranted to clarify the causal relationships among diet, HC-index, and human health.

The newly defined HC-index successfully positions the modern human gut microbiota within the broad dietary spectrum of mammals. While the human gut microbiota inherently retains herbivore-leaning potential, modern urbanized/industrialized lifestyles, including dietary shifts, have seemingly driven it toward a carnivore-leaning condition. Under modern lifestyles, the gut microbiota exists in a state of constant dietary fiber deficiency, which offers a mechanistic explanation for why fiber intake is so vital for human health. Indeed, higher HC-index values were positively associated with better health outcomes. The HC-index demonstrates enhanced applicability as a robust metric for comprehensively quantifying the diet–microbiota–health axis, thereby providing a novel framework for future human microbiota research.

## METHODS

### Study design and overview

This study integrated fecal gut microbiota datasets from diverse mammalian species and multiple human cohorts to develop and validate the HC-index, a quantitative microbiota-based metric representing the balance between herbivore- and carnivore-associated microbial signatures. Descriptions of each dataset, cohort-specific inclusion and exclusion criteria, sequencing methods, available metadata, and data accession numbers are detailed in Supplementary Method S1 and Supplementary Table S8.

### Animal fecal microbiota datasets

Fecal microbiota sequencing data from various mammalian species were compiled from the “SULAS Animal Dataset” (https://github.com/SULAS-H7/SULAS_anim_db), a collection of previously published and unpublished data generated by our laboratory, and two publicly available datasets^28,57^. Animals were classified as carnivores, herbivores, or omnivores based on diet categorizations in original studies; these were further checked using dietary information accessed from the Animal Diversity Web^58^ and EltonTraits database^19^. Nonfecal samples and those from nonmammalian hosts were excluded. The final dataset comprised 858 fecal samples from 109 mammalian species, including 343 from 35 carnivore species, 354 from 50 herbivores, and 161 from 24 omnivores. Sample-level accession information for animal fecal microbiota data is listed in Supplementary Table S9.

To minimize technical bias across studies, each dataset was processed independently utilizing QIIME 2 version 2024.5 with dataset-specific quality-filtering parameters. Taxonomy was assigned against the SILVA database version 138, and genus-level or genus-equivalent profiles were generated for downstream analyses. Potential dataset-specific effects were evaluated by PERMANOVA, including host diet and dataset identity as explanatory variables.

### Human fecal microbiota datasets

Human fecal microbiota profiles and associated metadata were compiled from seven cohorts or public datasets: the Human Microbiome Compendium dataset^22^, the J-MIC cohort (present study), the Kyotango cohort^24^, the KPUM dataset^26^, the American Gut Project^23^, the Hadza hunter-gatherer dataset^13^, and the dataset from the dietary fiber intervention study^27^.

The Human Microbiome Compendium was employed to evaluate the global applicability of the HC-index across adult populations and included 38,832 samples from seven geographic regions after filtering. The J-MIC cohort, comprising 26,783 adults, served as the primary cohort for ascertaining associations between the HC-index and questionnaire-derived dietary habits, lifestyle behaviors, stool characteristics, body mass index, and self-reported clinical histories. The Kyotango cohort included 770 adults with nutrient intakes estimated from the Brief-type self-administered Diet History Questionnaire^25^ and was used to examine associations between the HC-index and nutrient intake. The KPUM dataset included 1803 adults with comprehensive clinical blood test data and was utilized to assess associations with circulating clinical parameters. The American Gut Project included 3,678 adults and served as an external validation cohort for dietary associations in a non-Japanese population. The Hadza dataset included 158 samples and was used to position traditional human gut microbiota within the mammalian herbivore-carnivore microbial spectrum. The dietary fiber intervention study dataset was employed to evaluate the response of the HC-index to dietary intervention. This randomized controlled trial included 110 healthy adults who received dietary fiber supplementation for 4 weeks at either a low (2.2 g/day; *n* = 55) or a high (8.2 g/day; *n* = 55) dose. Cohort characteristics, inclusion and exclusion criteria, sample processing procedures, and dataset-specific filtering are detailed in Supplementary Method S1.

### Great ape fecal microbiota datasets

To compare HC-index values among urbanized humans, traditional populations, and the great ape species most closely related to humans, external fecal microbiota datasets from great apes were used^20,21^. Fecal microbiota profiles from chimpanzees (*n* = 38), bonobos (*n* = 11), gorillas (*n* = 40), and orangutans (*n* = 10) were analyzed to calculate HC-index values.

### Microbiota preprocessing and taxonomic profiling

Each cohort or dataset was processed independently using QIIME 2 version 2024.5 to minimize the influence of dataset-specific sequencing and preprocessing differences. Sequence reads were quality-filtered and denoised utilizing dataset-appropriate procedures, and taxonomy was assigned employing the SILVA database version 138. Genus-level or genus-equivalent relative abundance profiles were generated for downstream analyses. Dataset-specific processing workflows and analysis scripts are available in the associated github repository (https://github.com/SULAS-H7/HCI-study).

### Alpha and beta diversity analyses

Alpha diversity was assessed via Chao1 richness and Shannon diversity indices after rarefaction to 5,000 reads per sample; samples with fewer than 5,000 reads were excluded. Beta diversity was assessed using Bray-Curtis dissimilarity calculated from genus-level relative abundance profiles and visualized by principal coordinates analysis (PCoA). Effects of host diet, host taxonomic group, and dataset identity on microbial community composition were evaluated by PERMANOVA with 999 permutations.

### Identification of marker taxa for carnivores and herbivores

Marker taxa indicative of carnivore and herbivore fecal microbiota were identified using samples from animals classified as carnivores or herbivores. Taxa with a prevalence greater than 50% in either dietary group were retained as candidates, 61 from 963 detected taxa. Indicator species were analyzed using the indicspecies package in R, and P values were adjusted via the Benjamini–Hochberg procedure. Taxa satisfying adjusted *p* < 0.001 and predefined indicator-value thresholds were considered diet-associated. After excluding archaeal taxa due to their limited comparability across heterogeneous 16S rRNA amplification protocols^59^ as well as taxa unclassified at the genus or genus-equivalent level, the final marker set comprised four carnivore-associated taxa (C-taxa), including *Blautia* spp., *Escherichia–Shigella* spp., *Fusobacterium* spp., and *Peptoclostridium* spp., and 13 herbivore-associated taxa (H-taxa), including Bacteroidales p-251-o5, Christensenellaceae R-7 group, *Eubacterium coprostanoligenes* group, Oscillospiraceae NK4A214 group, Oscillospiraceae UCG-002, Oscillospiraceae UCG-005, Oscillospirales UCG-010, Prevotellaceae UCG-001, Prevotellaceae UCG-003, Prevotellaceae UCG-004, Rikenellaceae RC-9 group, *Ruminococcus* spp., and *Treponema* spp. Marker selection and threshold evaluation are detailed in the results section and Supplementary Table S2.

### Calculation of the herbivore-carnivore index

The HC-index was defined as the log-transformed ratio of the summed relative abundance of H-taxa to that of C-taxa. For each sample, the HC-index was calculated as,

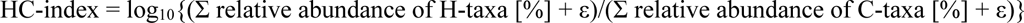

where ε denotes the pseudo-count added to both the numerator and denominator to avoid division by zero. As alpha diversity was analyzed after rarefaction to 5,000 reads per sample, the detection limit was set at 0.01%, corresponding to approximately one read per 5,000–10,000. The pseudo-count ε was set to 0.005%, corresponding to half the detection limit.

Higher HC-index values indicate a greater relative abundance of herbivore-associated taxa and, therefore, a more herbivore-leaning gut microbiota profile. Lower HC-index values indicate a greater relative proportion of carnivore-associated taxa and, therefore, a more carnivore-leaning profile.

As an exploratory analysis, we also constructed a simplified HC-index using seven selected marker taxa instead of the full 17-taxon marker set: *Ruminococcus* spp., Oscillospiraceae UCG-002, Christensenellaceae R-7 group, *Eubacterium coprostanoligenes* group, *Blautia* spp., *Fusobacterium* spp., and *Escherichia–Shigella* spp. The performance of this simplified HC-index was evaluated relative to the full index (Supplementary Fig. S9).

### Validation of the HC-index in animal and human datasets

The HC-index was validated by comparing its distribution among carnivores, herbivores, and omnivores using Kruskal–Wallis tests followed by pairwise Wilcoxon rank-sum tests with Benjamini–Hochberg correction. Its association with the proportion of plant-based foods in host diets was evaluated using dietary information obtained from the EltonTraits database^19^ by Spearman rank correlation. Classification performance in herbivore and carnivore samples was assessed employing an HC-index threshold of 0: samples with values ≥ 0 were classified as herbivore-like, and those with values < 0 as carnivore-like.

Associations between the HC-index, alpha diversity indices, and PCoA coordinates across animal samples and within individual dietary groups were evaluated utilizing Spearman rank correlation. The HC-index was subsequently applied to urbanized human, Hadza, and nonhuman ape gut microbiota datasets to position these populations within the mammalian herbivore-carnivore microbial spectrum.

### Dietary, lifestyle, and clinical association analyses

In the J-MIC cohort, associations between HC-index and questionnaire-derived variables were evaluated using regression models adjusted for age and sex. Multicategory outcomes, including dietary intake frequencies, lifestyle behaviors, and stool characteristics, were analyzed via multinomial logistic regression, whereas binary outcomes, including self-reported clinical histories, were examined employing binary logistic regression. In all models, HC-index was included as the main explanatory variable, with age and sex as covariates. Adjusted predicted probabilities were estimated from fitted models for visualization across the observed HC-index range. Model specifications are detailed in Supplementary Method S2.

In the Kyotango cohort, associations between HC-index and nutrient intake variables derived from the Brief-type self-administered Diet History Questionnaire were evaluated using linear regression models adjusted for age and sex. Both absolute and energy-adjusted nutrient intakes were examined, and P values were adjusted for multiple testing via the Benjamini–Hochberg procedure.

In the KPUM dataset, associations between HC-index and clinical blood parameters were evaluated employing linear regression models adjusted for age and sex. P values were adjusted for multiple testing via the Benjamini–Hochberg procedure.

### Comparison of explanatory performance between indices

To determine whether the HC-index captured information distinct from conventional alpha diversity measures, its associations and explanatory performance were compared with those of Chao1 richness and Shannon diversity. Age- and sex-adjusted relations between the HC and alpha diversity indices were ascertained using Spearman correlation after residualization against age and sex.

For categorical outcomes, separate age- and sex-adjusted models were fitted utilizing the HC-index or an alpha diversity index as the main explanatory variable, and model improvement relative to covariate-only models was quantified employing likelihood-ratio statistics. For continuous outcomes, explanatory performance was compared based on the incremental adjusted *R²* contributed by each microbiota metric beyond age and sex. Statistical significance of model improvement was assessed via likelihood-ratio or partial F-tests, as appropriate, with Benjamini–Hochberg correction. Definitions of the comparative metrics are detailed in Supplementary Method S3.

### Shotgun metagenomic sequencing and functional analysis

Shotgun metagenomic sequencing used fecal DNA from 156 randomly selected J-MIC participants. Libraries were prepared from 200 ng of DNA using the NEBNext Ultra II FS DNA Library Prep Kit for Illumina and sequenced on the NovaSeq X platform. Sequencing data were processed using the default pipeline of SqueezeMeta^60^ v1.7.0 in co-assembly mode. Briefly, reads were assembled with MEGAHIT^61^, open reading frames were predicted with Prodigal^62^, and taxonomic annotation against the NCBI nonredundant protein database used the LCA-based procedure implemented in SqueezeMeta. Functional annotation was against the KEGG database and a custom dbCAN3-derived CAZyme database. Gene abundances were quantified as transcripts per million and aggregated at the KEGG Orthology and CAZyme-family levels.

Associations between HC-index and metagenomic functional profiles were evaluated using Bray-Curtis dissimilarity and PCoA. To identify HC-index-associated CAZyme families, samples in the lowest and highest HC-index quartiles were compared via Wilcoxon rank-sum tests with Benjamini–Hochberg correction. Significant glycoside hydrolase families were categorized according to predicted substrate classes based on Smits et al. (2017). Filtering criteria and enrichment-score calculations are detailed in Supplementary Method S4.

### Statistical analysis

All statistical analyses were performed in R version 4.6.1. Unless otherwise specified, all tests were two-sided. When multiple hypotheses were examined, P values were adjusted using the Benjamini–Hochberg procedure. Statistical procedures specific to individual analyses are described in the corresponding Methods sections.

### Ethics approval and participant consent

The J-MIC cohort, Kyotango cohort, and KPUM dataset were generated by our group and were approved by the ethics committee of Kyoto Prefectural University of Medicine under approval numbers ERB- C-2182-2, ERB-C-885-10, and ERB-C-534-10, respectively. All participants provided written informed consent. For the remaining publicly available datasets, ethical approval and informed consent were obtained in the original studies.

## ACKNOWLEDGEMENTS

The authors sincerely thank Mai Sakagami and Megumi Watanabe of Yokohama Zoological Gardens and Daisuke Nakagawa and Sachiko Ezaki of Kobe Animal Kingdom for their assistance with data collection.

## AUTHOR CONTRIBUTIONS

Conceptualization, H.M. and R.I.

Methodology, H.M., To.Ta., Ta.Tsu. and R.I.

Software, H.M. and R.I.

Validation, H.M. and R.I.

Formal analysis, H.M. and R.I.

Investigation, H.M., To.Ta., S.I. and R.I.

Resources, To.Ta., D.U., A.K., M.O., H.I., K.N., H.K., S.M., Y.N. and R.I.

Data curation, H.M., To.Ta., D.U., A.K., M.O., S.I., H.K., S.M. and R.I.

Writing—original draft, H.M. and R.I.

Writing—review and editing, To.Ta., Ta.Tsu., Y.N. and R.I.

Visualization, H.M. and R.I.

Supervision, R.I.

Project administration, R.I.

## DATA AVAILABILITY

Raw sequencing data for each study can be accessed as described in Supplementary Table S8. The genus-level composition table is available at the github repository (https://github.com/SULAS-H7/HCI-study). Metadata of the J-MIC, Kyotango, and KPUM cohort/datasets containing clinical and nutritional backgrounds cannot be deposited publicly due to participant privacy and ethical restrictions. Access to this metadata can be requested by contacting the corresponding author, subject to institutional ethical compliance. All other data supporting the findings of this study are available in this article and its Supplementary Information files, or from the corresponding author on request.

## CODE AVAILABILITY

The analytical code used to generate the results is publicly available on the github repository (https://github.com/SULAS-H7/HCI-study).

---

**Fig. S1.**
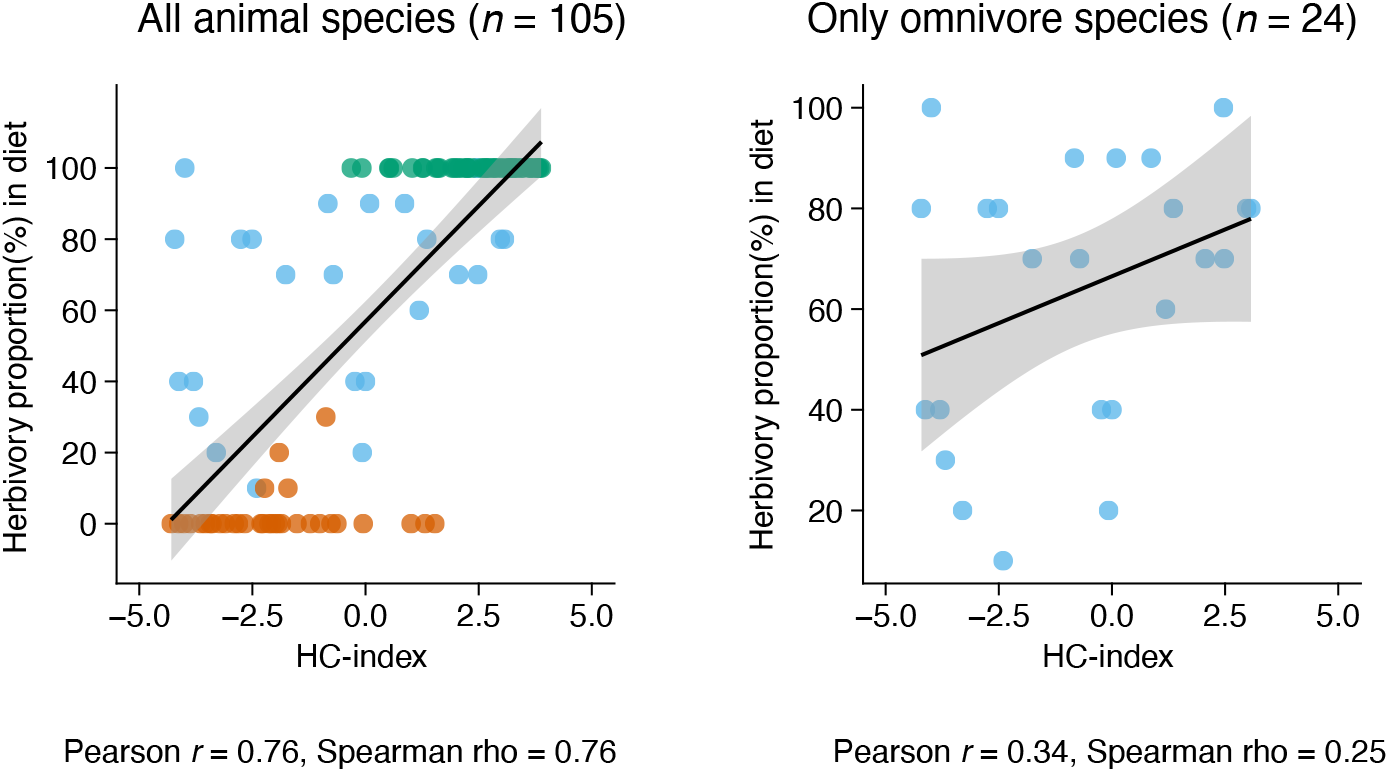
Correlation between the HC index and the proportion of plant-based foods in the diet across diverse animal species. HC-index values were averaged within each animal species. The proportion of plant-based foods in the diet of each animal species was calculated using the EltonTraits database (Wilman et al., 2014). Four animal species without dietary information in the EltonTraits database were excluded from this analysis. A total of 105 animal species, including 33 carnivores (red dots), 48 herbivores (green dots) and 24 omnivores (blue dots), were included. The black line and grey ribbon represent the regression line and its 95% confidence interval, respectively. Correlation coefficients were calculated as both Pearson’s *r* and Spearman’s rho.

**Fig. S2.**
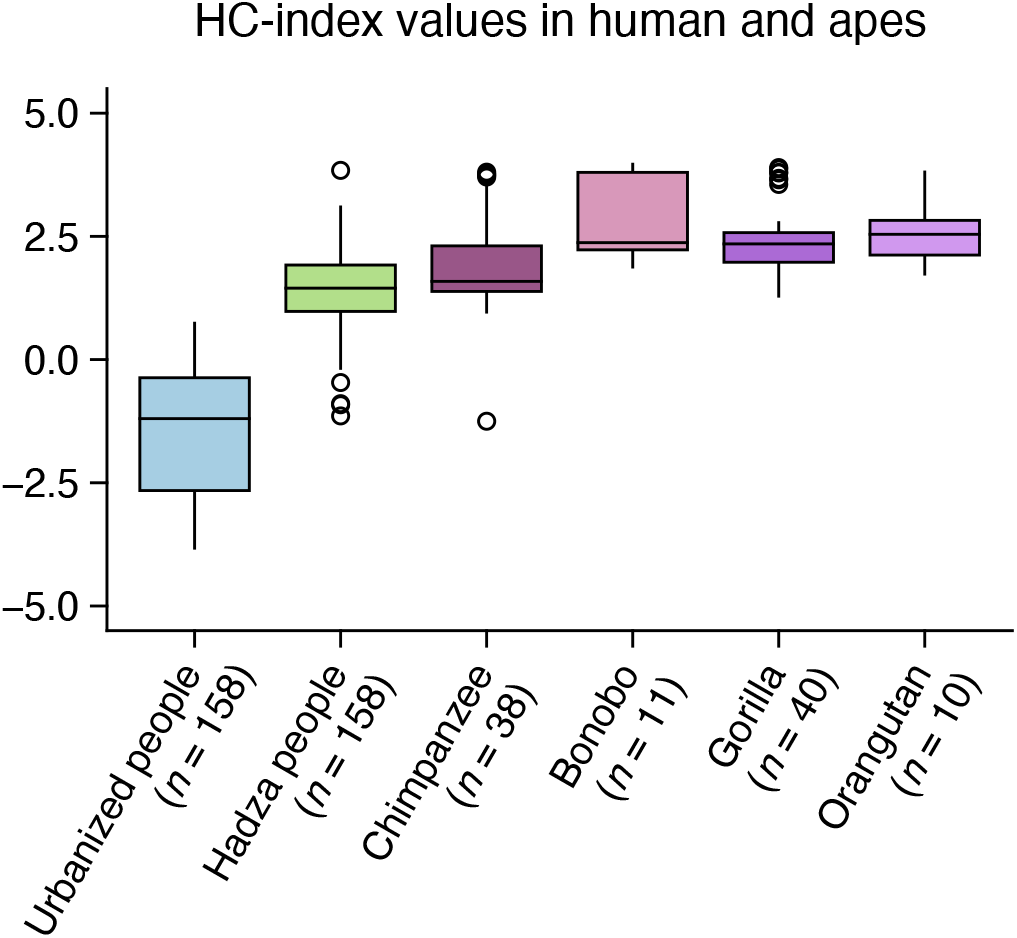
Comparison of HC index values between human populations and non-human great apes. Urbanized human and Hadza data were randomly selected from the J-MIC cohort and Hadza cohort datasets, respectively. Data from non-human great apes, including chimpanzees, bonobos, gorillas and orangutans, were obtained from Narat et al. (2020) and Nishida et al. (2021) and pooled. Details of the data analysis are described in Supplementary Method S1.

**Fig. S3.**
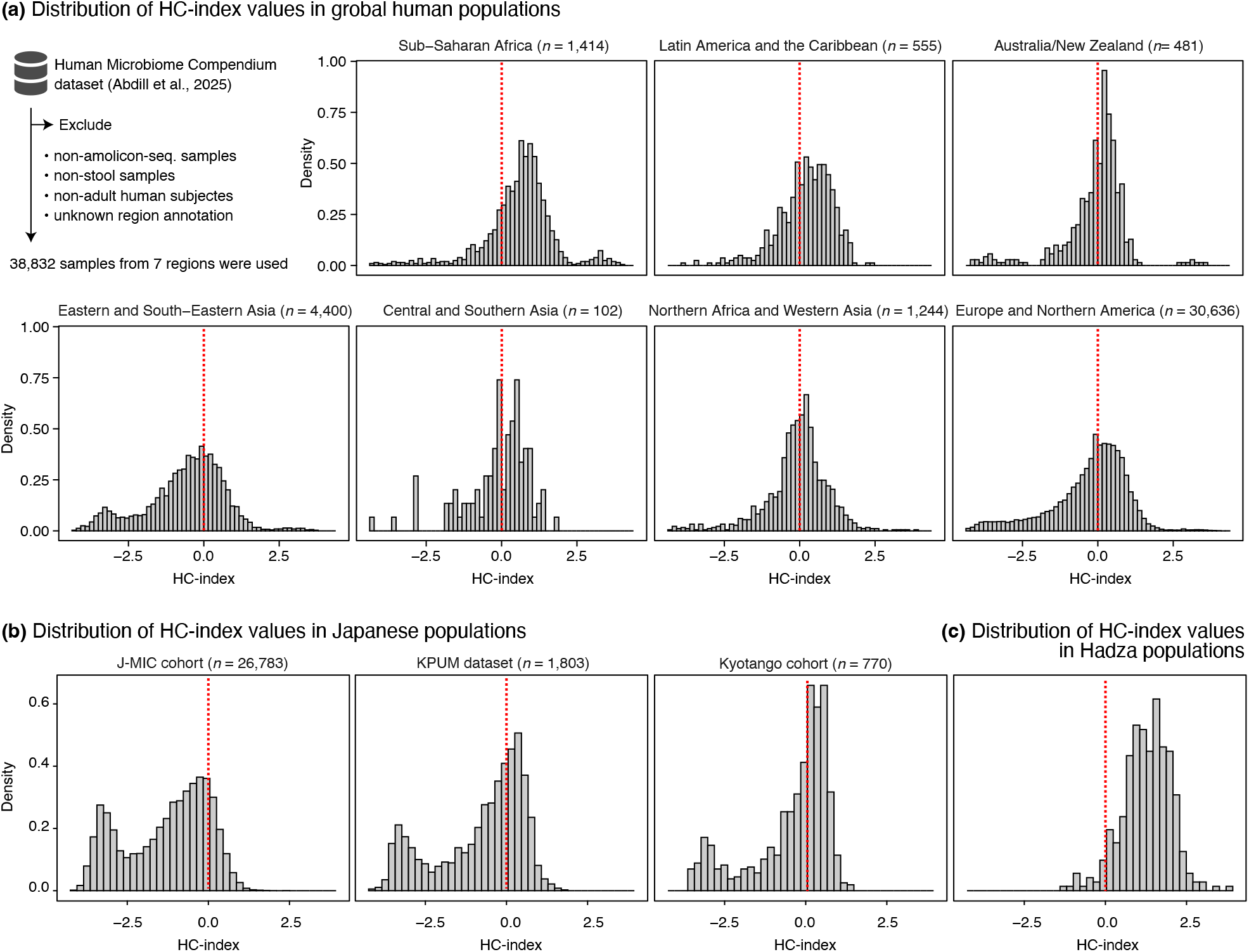
Distribution of HC index values across human populations from different geographic regions. Distribution of HC index values in **(a)** global human populations from different geographic regions in the Human Microbiome Compendium study, **(b)** Japanese populations from our studies and **(c)** the Hadza population. Details of the data analysis are described in Supplementary Method S1.

**Fig. S4.**
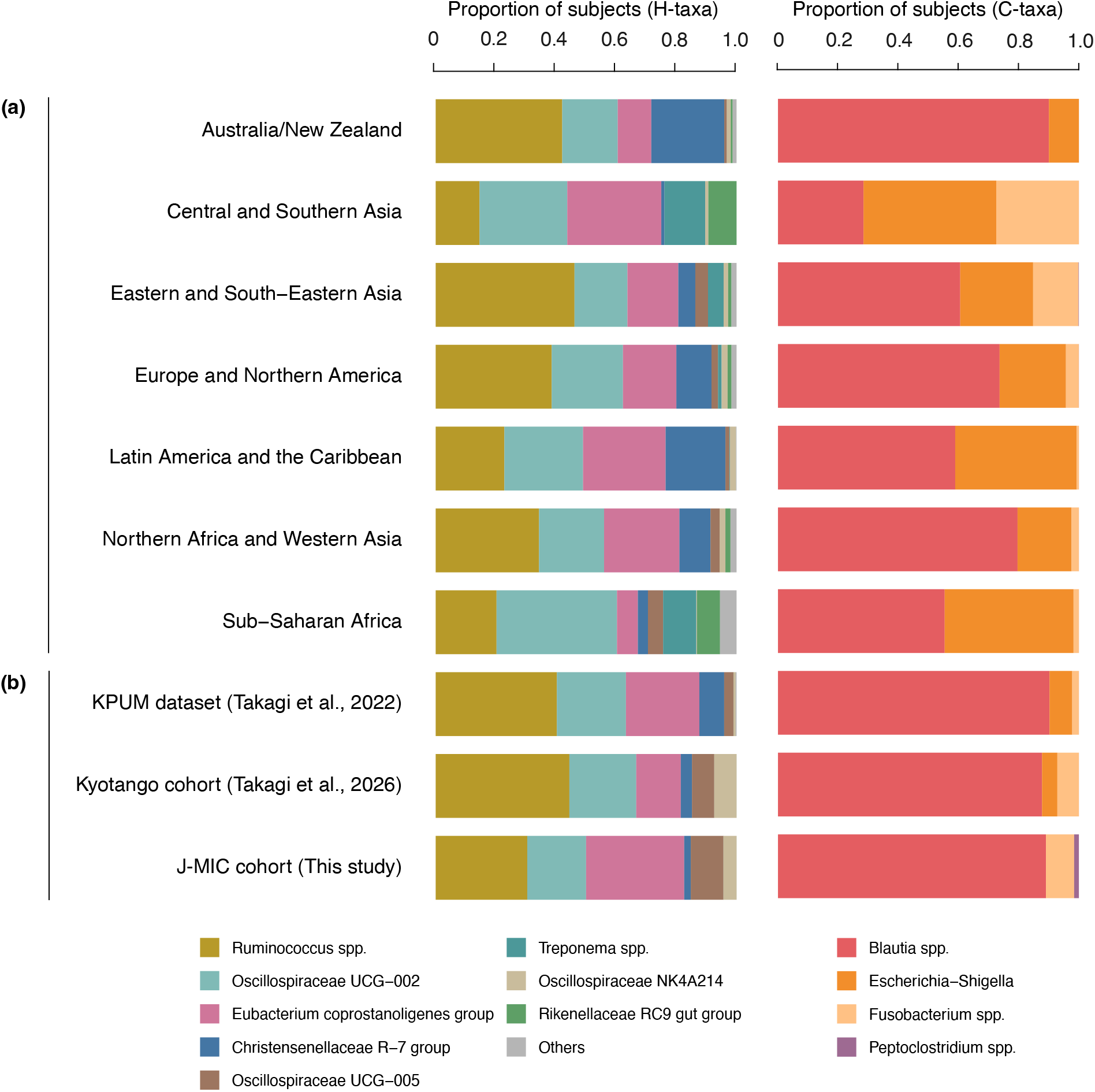
Proportions of subjects by predominant H- and C-taxa across human populations from different geographic regions. The stacked bars represent the proportions of subjects classified according to their most abundant taxon within the H-taxa and C-taxa, respectively. Data are shown for **(a)** human populations from different geographic regions included in the Human Microbiome Compendium study and **(b)** Japanese populations from our studies.

**Fig. S5.**
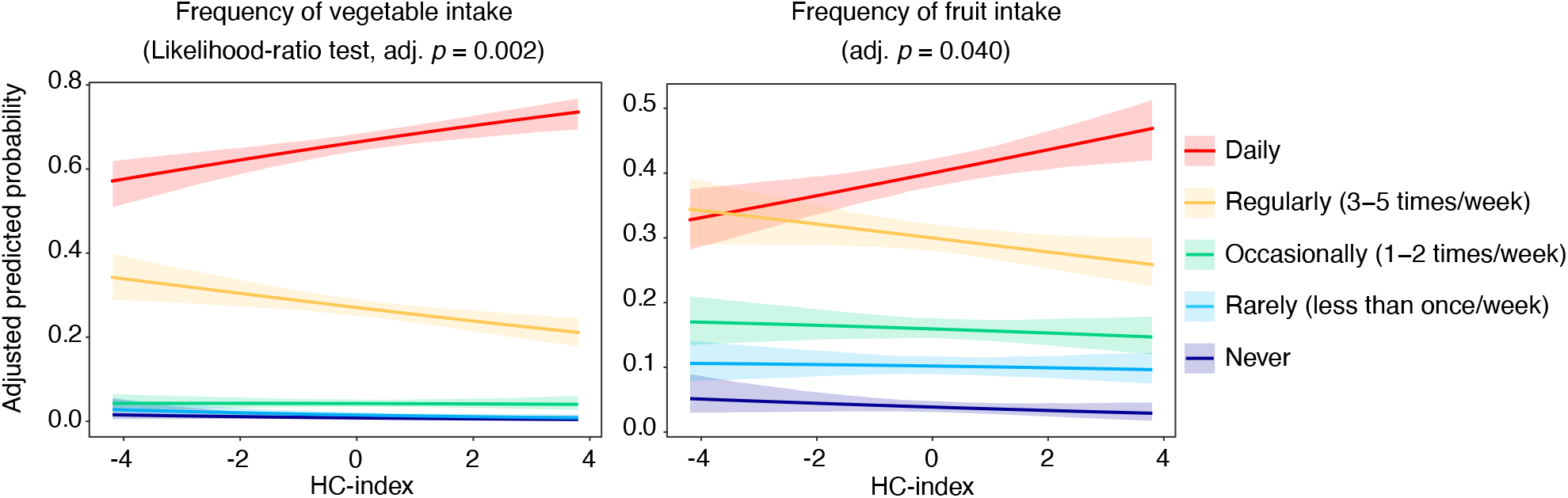
Associations between the HC index and questionnaire-derived frequency of vegetable and fruit intake in the American Gut Project dataset. Age- and sex-adjusted multinomial logistic regression models were used to estimate predicted marginal probabilities for each response category. The curves and shaded areas represent the adjusted predicted probabilities and their 95% confidence intervals, respectively. Overall statistical significance of the HC-index was evaluated using a likelihood-ratio test comparing a full model including the HC-index, age, and sex with a reduced model comprising only age and sex.

**Fig. S6.**
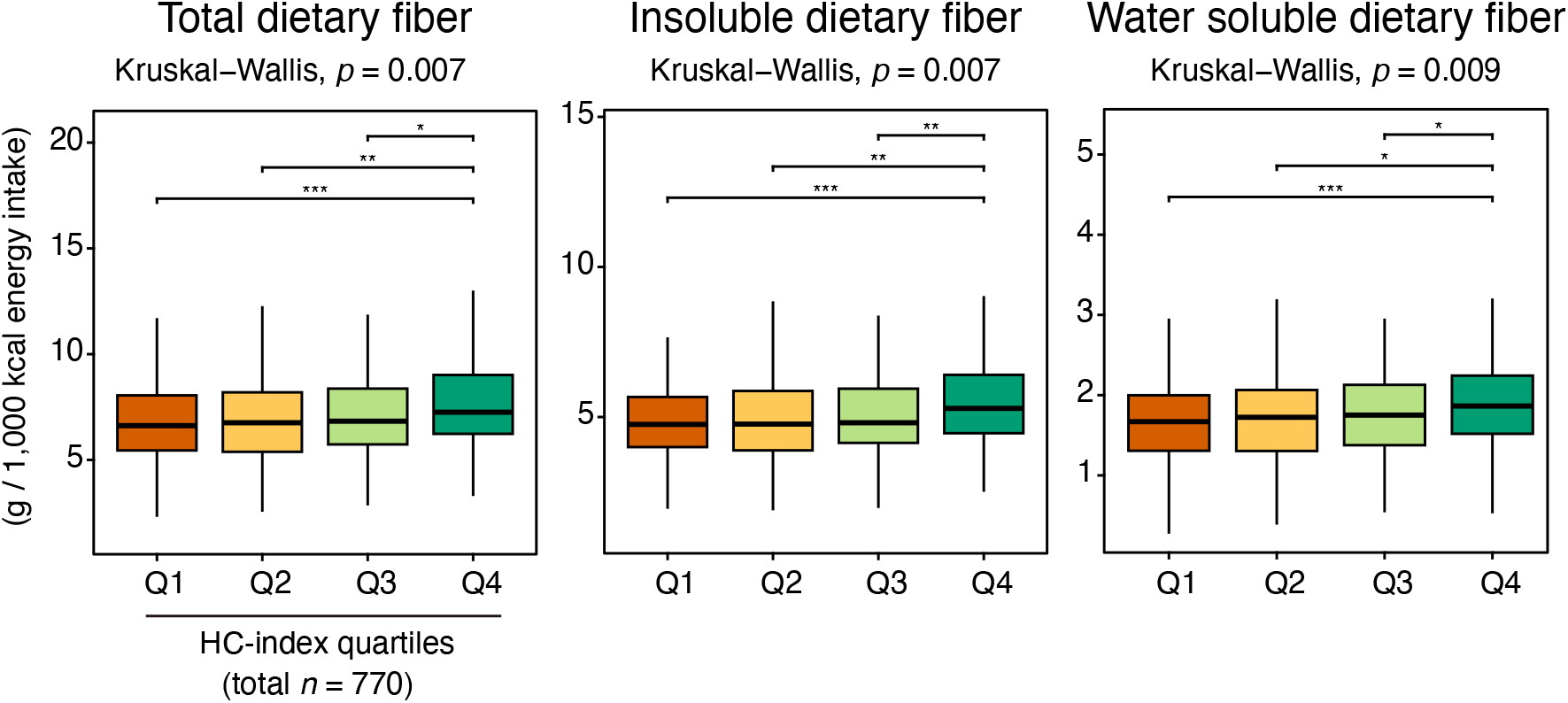
Comparison of dietary fiber intake across HC index quartiles in the Kyotango cohort. Energy-adjusted intakes of total, insoluble, and water-soluble dietary fiber were compared across HC index quartiles. Nutrient intake was estimated using the Brief-type self-administered Diet History Questionnaire (BDHQ), and dietary fiber intake was adjusted per 1,000 kcal of energy intake. Statistical significance was assessed using a Kruskal–Wallis test followed by two-sided pairwise Wilcoxon rank-sum tests with Benjamini–Hochberg correction. Significance levels are indicated as *adjusted *p* < 0.05, **adjusted *p* < 0.01 and ***adjusted *p* < 0.001.

**Fig. S7.**
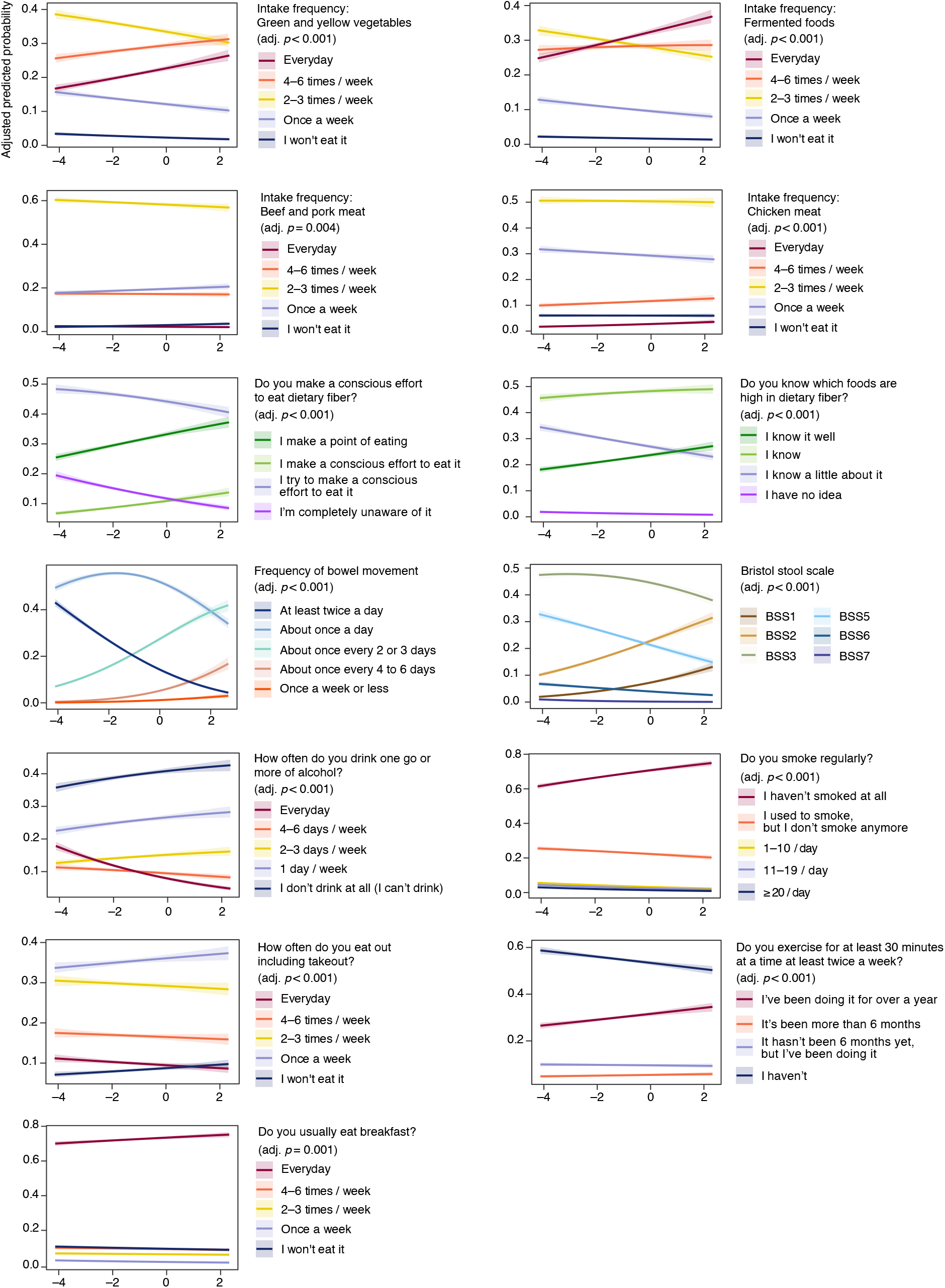
**Associations between the HC index and the questionnaire-derived dietary habits, stool characteristics, and lifestyle behaviors in the J-MIC cohort**. Age- and sex-adjusted multinomial logistic regression models were used to estimate predicted probabilities for each response category across the observed range of HC-index values. The curves and shaded areas represent the adjusted predicted probabilities and their 95% confidence intervals, respectively. Statistical significance of the HC-index was evaluated using a likelihood-ratio test comparing a full model including the HC-index, age, and sex with a reduced model comprising only age and sex. P values were adjusted for multiple testing by applying the Benjamini–Hochberg procedure.

**Fig. S8.**
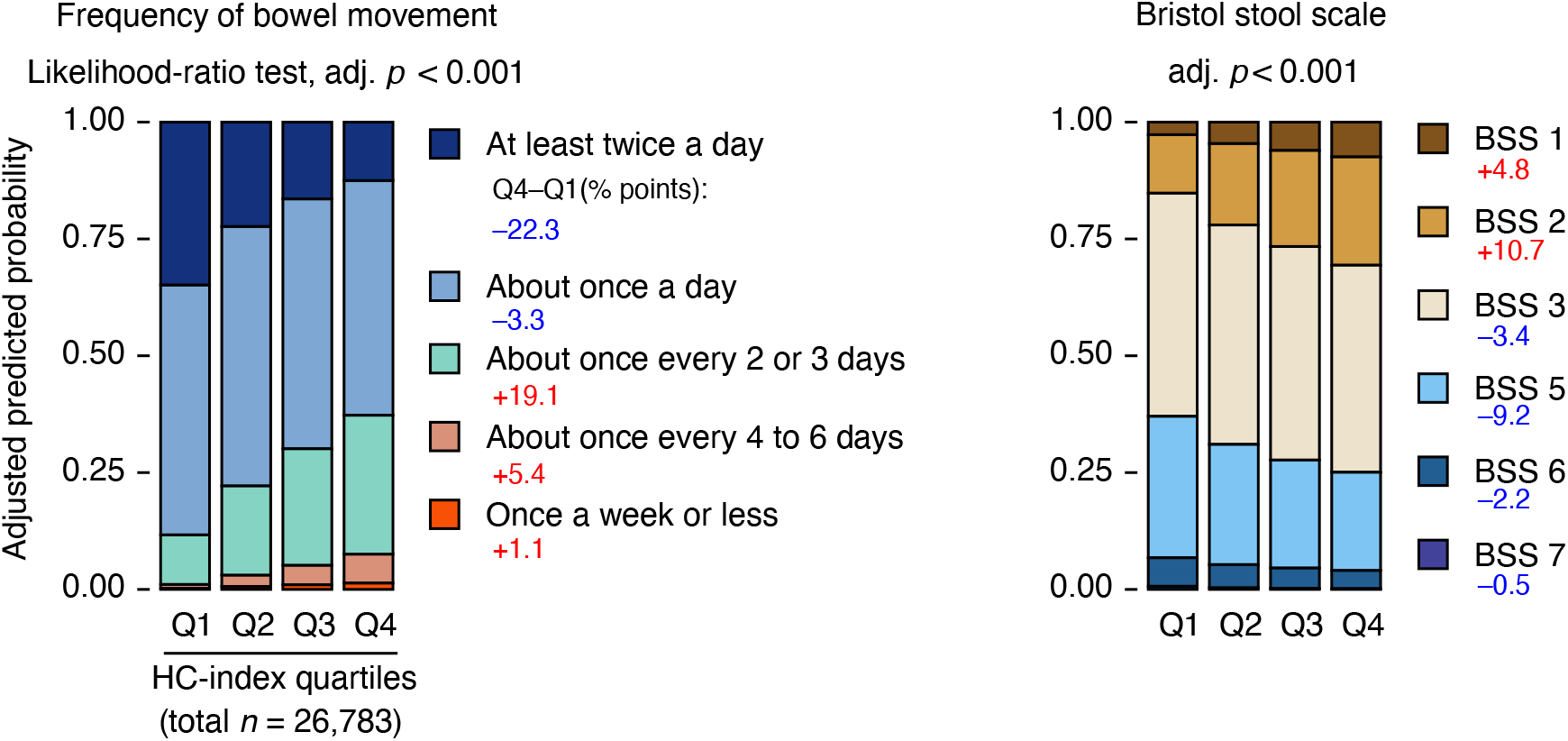
Associations between the HC index and questionnaire-derived bowel movement frequency and stool conditions in the J-MIC cohort. Age- and sex-adjusted multinomial logistic regression models were used to predict marginal probabilities for each response category, shown as stacked bars across HC index quartiles. Statistical significance of the HC-index was evaluated using a likelihood-ratio test comparing a full model including the HC-index, age, and sex with a reduced model comprising only age and sex. P values were adjusted for multiple testing by applying the Benjamini– Hochberg procedure. For interpretation, Q4−Q1 differences in adjusted predicted probabilities are shown next to the stacked bars in percentage points; positive values indicate a higher predicted probability in Q4 (higher HC-index), whereas negative values indicate a higher predicted probability in Q1 (lower HC-index). The corresponding model-based curves are presented in Supplementary Fig. S7.

**Fig. S9.**
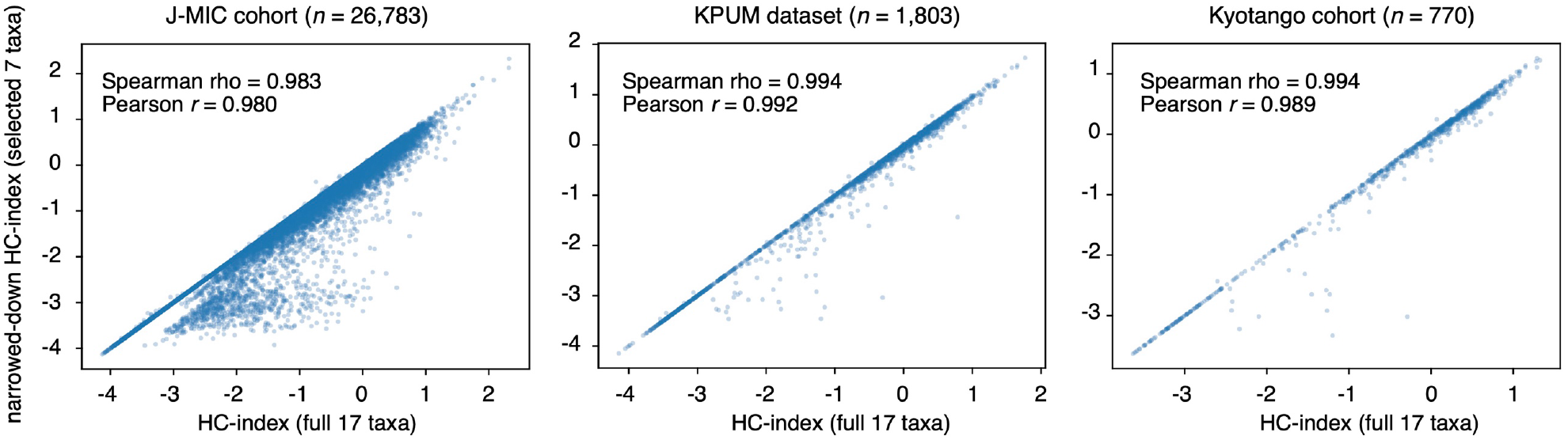
Correlation between the full HC index and a simplified HC index. As an exploratory analysis, a simplified HC index was constructed using 7 selected marker taxa instead of the full 17-taxon marker set: 4 selected herbivore-associated taxa, including *Ruminococcus* spp., Oscillospiraceae UCG-002, Christensenellaceae R-7 group and *Eubacterium coprostanoligenes* group, and 3 selected carnivore-associated taxa, including *Blautia* spp., *Fusobacterium* spp. and *Escherichia–Shigella* spp. The performance of the simplified HC index was evaluated in Japanese populations from our studies by comparing it with the full HC index based on all 17 marker taxa. Pearson’s r and Spearman’s rho were calculated to assess the correlation between the two indices.

## Supplementary Method S1. Detailed descriptions of human datasets and cohorts and dataset-specific processing

### Human Microbiome Compendium dataset

The Human Microbiome Compendium dataset (Abdill et al., 2025) was used to establish the global applicability of the HC-index across adult human populations. After filtering by metadata and analysis-specific inclusion criteria, 38,832 adult stool samples from 7 geographic regions were included. The taxonomic abundance table was downloaded and converted from taxon counts to per-sample relative abundances. The HC-index was then calculated as described in the Methods section of the main text and merged with sample- and project-level metadata. Downstream analyses were restricted to stool samples from adult human subjects profiled via amplicon sequencing. Adult human samples were identified by keyword-based filtering of the subject-description field, excluding non-human samples and those from infants, children, adolescents, or mixed-age cohorts. To reduce the possibility of technical artifacts, projects were excluded if ≥ 20% of their samples had near-zero HC-index values, defined as |HC-index| < 0.01. For geographic analyses, samples with unknown regional annotations were excluded, as were regions represented by < 30 samples. After these filtering procedures, 38,832 samples from seven geographic regions were retained.

### J-MIC cohort

The J-MIC cohort served as the primary human cohort to evaluate associations between HC-index and questionnaire-derived dietary habits, lifestyle behaviors, stool characteristics, body mass index, and self-reported clinical histories. The cohort comprised 26,783 adults with standardized fecal microbiota profiles and questionnaire-based background information. Fecal sample collection, DNA extraction, and 16S rRNA gene amplicon sequencing were according to described procedures (Takagi et al., 2022). For comparisons involving animal samples, urbanized human populations, and traditional populations, J-MIC samples were co-analyzed with animal and other human fecal microbiota profiles post-harmonization at the genus or genus-equivalent levels (Fig. 1d). For principal coordinate analysis (PCoA) visualization, 30 J-MIC samples were randomly selected to reduce imbalance in sample size among compared groups. In addition, for comparing HC-index values among urbanized human populations, traditional populations, and great apes (Supplementary Fig. S2), J-MIC samples served as representative urbanized human samples. To match the sample size of the Hadza cohort, 158 J-MIC samples were randomly selected.

### KPUM dataset

The KPUM dataset (Takagi et al., 2022) was used to evaluate associations between HC-index and clinical blood parameters. The dataset comprised 1,803 adults of both sexes, including 283 individuals classified as healthy, with fecal microbiota profiles and clinical blood test data. Detailed participant enrollment procedures, cohort characteristics, fecal sample collection, DNA extraction, and 16S rRNA gene amplicon sequencing were as described originally (Takagi et al., 2022).

### Kyotango cohort

The Kyotango cohort (Takagi et al., 2026) served as an independent cohort with detailed estimates of nutrient intake. The cohort comprised 770 adults with dietary intake data derived from the Brief-type self-administered Diet History Questionnaire (BDHQ; Kobayashi et al., 2012) and fecal microbiota data. Detailed participant enrollment procedures, cohort characteristics, fecal sample collection, DNA extraction, and 16S rRNA gene amplicon sequencing were as described originally (Takagi et al., 2026). Nutrient intake variables employed in the present analyses included both absolute intake and energy-adjusted values expressed per 1,000 kcal, as described in the corresponding procedures for statistical analysis.

### American Gut Project dataset

The American Gut Project dataset (McDonald et al., 2018) served as an independent external validation cohort to examine whether associations between HC-index and dietary habits were reproducible in a non-Japanese population. Metadata for Qiita study 10317 was obtained from Qiita. Empty strings and the entries “Not provided,” “not provided,” “Unspecified,” and “unspecified” were treated as missing values; duplicate “sample_name” entries were removed. For diet-related analyses, samples were retained when data were available for “sample_name,” “diet_type,” “red_meat_frequency,” “vegetable_frequency,” “meat_eggs_frequency,” and “fruit_frequency.” To reduce imbalance among dietary categories, up to 1,000 samples per “*diet_type*” category were randomly selected utilizing a fixed random seed. This resulted in a final dataset of 3,678 adults. Microbiome feature tables corresponding to the selected sample identifiers were retrieved from redbiom using the Deblur 2021.09 Illumina 16S V4 contexts for both 90-nt (“Deblur_2021.09-Illumina-16S-V4-90nt-dd6875”) and 125-nt (“Deblur_2021.09-Illumina-16S-V4-125nt-92f954”) sequences, with ambiguity resolution set to “most-reads.” Taxonomic profiles were generated in QIIME 2 version 2024.5 and the SILVA 138 reference database. The 125-nt dataset was employed preferentially, and samples available only in the 90-nt dataset were added after harmonization at the genus or genus-equivalent level. Samples in which a single genus accounted for > 80% of the total relative abundance were excluded. Detailed scripts for sample selection, redbiom data retrieval, taxonomic assignment, integration of the 90-nt and 125-nt datasets, and downstream processing are available in the related github repository (https://github.com/SULAS-H7/HCI-study).

### Hadza dataset

The Hadza dataset (Smits et al., 2017) was primarily used to position traditional human gut microbiota within the mammalian herbivore–carnivore microbial spectrum and to compare HC-index distributions among urbanized humans, Hadza individuals, and non-human apes. Calculation of the HC-index and subsequent analyses were restricted to stool samples from adult human participants (aged ≥20 years; *n* = 158). For cross-dataset comparisons, Hadza samples were co-analyzed with animal and urbanized human fecal microbiota profiles post-harmonization at the genus or genus-equivalent level. For PCoA visualization, 30 Hadza samples were randomly selected using a fixed random seed to reduce imbalance in sample size among comparison groups. In addition, for comparing HC-index values among urbanized human populations, traditional populations, and great apes (Supplementary Fig. S2), Hadza samples were served as representative traditional human samples.

## Supplementary Method S2. Detailed model specifications

### General regression framework

Associations between HC-index and human phenotypic variables were evaluated via regression models adjusted for age and sex. In all models, the HC-index was included as the main explanatory variable, with age and sex as covariates. The model choice depended on the scale and structure of each outcome variable. Multicategorical outcomes were analyzed using multinomial logistic regression, binary outcomes utilizing binary logistic regression, and continuous outcomes employing linear regression. Unless otherwise specified, statistical significance of the HC-index was assessed using the model-specific test described below, and P-values were adjusted for multiple testing by applying the Benjamini–Hochberg procedure.

### Categorical dietary, lifestyle, and stool-related outcomes in the J-MIC cohort

Questionnaire-derived dietary habits, lifestyle behaviors, and stool characteristics were analyzed as multi-categorical outcomes via multinomial logistic regression models. For each outcome, the overall statistical significance of the HC-index was evaluated using a likelihood-ratio test comparing a full model including the HC-index, age, and sex with a reduced model comprising only age and sex. Within each response variable, P values were adjusted for multiple testing utilizing the Benjamini–Hochberg procedure. For interpretation, adjusted predicted probabilities were estimated from the fitted models across the range of HC-index values observed. Age was fixed at the median age of the participants, and predicted probabilities were averaged across sex strata. For multinomial logistic regression models, 95% confidence intervals for predicted category probabilities were estimated by simulation from the fitted coefficient covariance matrix.

For additional descriptive summaries, predicted probabilities were also estimated across the four HC-index quartiles using the median HC-index value within each quartile. Post-hoc pairwise comparisons among quartiles were not employed as the primary inferential procedure. Instead, Q4-minus-Q1 contrasts in adjusted predicted probabilities were calculated for multinomial outcomes utilizing simulation-based estimates and were applied for descriptively interpreting the effect magnitude.

### Binary clinical history outcomes in the J-MIC cohort

Self-reported clinical histories were treated as binary outcomes and analyzed via binary logistic regression models adjusted for age and sex. Overall significance was evaluated by comparing the full model including the HC-index, age, and sex with the corresponding reduced model only including age and sex using a likelihood-ratio test. Odds ratios were calculated per one-unit increase in the HC-index, with Wald-type 95% confidence intervals derived from the estimated regression coefficient and its standard error. P values were adjusted for multiple testing by applying the Benjamini–Hochberg procedure. Disease-related outcomes meeting the pre-specified BH-adjusted significance threshold were additionally summarized in forest plots.

For visualizing binary outcomes, adjusted predicted probabilities were estimated across the HC-index range observed with age fixed at the median age of the participants, and predictions averaged across sex strata. Approximate 95% confidence intervals were calculated on the link scale and back-transformed to the probability scale.

### Body mass index in the J-MIC cohort

Body mass index (BMI) was analyzed as a continuous outcome via linear regression models adjusted for age and sex. Participants with BMI values outside 15–40 kg/m² were excluded. The regression coefficient for the HC-index represented the adjusted change in BMI associated with a one-unit increase in HC-index. Statistical significance was assessed using the Wald test statistic for the HC-index coefficient. P values were adjusted for multiple testing by employing the Benjamini–Hochberg procedure where applicable. For stratified visualization, BMI distributions across HC-index quartiles were examined separately by sex among participants aged 40–60 years.

### Nutrient intake analyses in the Kyotango cohort

Nutrient intakes estimated using the Brief-type self-administered Diet History Questionnaire (BDHQ; Kobayashi et al., 2012) were analyzed as continuous outcomes, and partial Pearson correlation coefficients between each nutrient intake variable and the HC-index and P values were calculated post-adjustment for age and sex. Both absolute and energy-adjusted nutrient intakes expressed per 1,000 kcal were analyzed. P values were adjusted for multiple testing by applying the Benjamini–Hochberg procedure.

### Clinical blood parameter analyses in the KPUM dataset

Clinical blood parameters were analyzed as continuous outcomes, and partial Pearson correlation coefficients between each clinical blood parameter and the HC-index and P values were calculated post-adjustment for age and sex. P values were adjusted for multiple testing by applying the Benjamini–Hochberg procedure.

## Supplementary Method S3. Comparison of explanatory performance between the HC-index and alpha diversity indices

To determine whether the HC-index captured information distinct from conventional alpha diversity measures, its relationships with and explanatory performance relative to Chao1 richness and Shannon diversity were evaluated.

First, associations between the HC-index and each alpha diversity index were assessed post-adjustment for age and sex. They were then separately residualized with respect to age and sex via linear regression models, and Spearman rank correlations were then calculated between the residuals obtained. P values were adjusted by applying the Benjamini–Hochberg procedure.

For categorical outcomes, explanatory performance was compared via age- and sex-adjusted regression models fitted separately with either the HC or an alpha diversity index as the main explanatory variable. Binary category outcomes were analyzed using logistic regression models, whereas multicategory outcomes were analyzed utilizing multinomial logistic regression models. For each model, improvement over a covariate-only model including age and sex was quantified employing the likelihood-ratio test statistic (LRT). Differences in explanatory performance were summarized as,

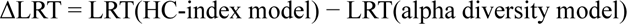

Where the alpha diversity model included either Chao1 richness or Shannon diversity. Positive ΔLRT values indicated greater model improvement for the HC-index, whereas negative values suggested the contrary.

For continuous outcomes, explanatory performance was compared via age- and sex-adjusted linear regression models. For each outcome, three models were fitted: a covariate-only model including age and sex, a model adding the HC-index, and a model additionally comprising an alpha diversity index. Model improvement was quantified based on adjusted *R²*. The additional explanatory contribution of each microbiota metric relative to the covariate-only model was calculated, and differences in explanatory performance were summarized as,

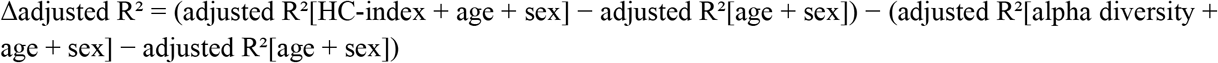

Positive Δadjusted *R²* values indicated a greater additional explanatory contribution of the HC-index, whereas negative values suggested the opposite.

Statistical significance of model improvement for continuous outcomes was evaluated using partial *F*-tests comparing each full model with the corresponding covariate-only model. P values were adjusted by applying the Benjamini–Hochberg procedure separately for comparisons involving each alpha diversity metric.

## Supplementary Method S4. Detailed methodology of shotgun metagenomic sequencing analysis

To identify CAZyme families associated with low and high HC-index values, samples were divided into HC-index quartiles, and those in the lowest quartile (Q1) and highest quartile (Q4) were compared. CAZyme features belonging to glycoside hydrolase (GH), polysaccharide lyase (PL), carbohydrate-binding module (CBM), and carbohydrate esterase (CE) classes were analyzed. Features detected in ≥ 10% of the Q1/Q4 samples were retained. Q1 and Q4 abundances were compared using the Wilcoxon rank-sum test, and P values were adjusted using the Benjamini-Hochberg procedure. Features with adjusted P values < 0.05 were considered significant. For such features, LDA scores were calculated using linear discriminant analysis applied to log1p-transformed transcripts per million (TPM) values, and enrichment direction was assigned according to the higher mean TPM in Q1 or Q4. Positive signed LDA scores indicate Q4 enrichment, whereas negative signed LDA scores indicate Q1 enrichment. GH families were categorized by predicted substrate class as targeting plant cell wall carbohydrates, animal-derived carbohydrates, mucin-related glycans, or other substrates (Smits et al., 2017).

